# Preserved particulate organic carbon is likely derived from the subsurface sulfidic photic zone of the Proterozoic Ocean: evidence from a modern, oxygen-deficient lake

**DOI:** 10.1101/2023.11.19.567733

**Authors:** Ashley Brooke Cohen, Gordon T Taylor, Gregory Henkes, Felix Weber, Lisa N Christensen, Milana Yagudaeva, Evan Lo, Michael McCormick

**Affiliations:** U. S. Naval Research Laboratory, Washington, DC, USA; Stony Brook University, NY, USA; Alfred-Wegener-Institut Helmholtz-Zentrum, Helgoland, Germany; Cold Spring Harbor Laboratory, NY, USA; Moderna; Hamilton College

**Keywords:** carbon fixation pathways, meromictic, carbon stable isotopes, particle-associated, Fayetteville Green Lake, anoxygenic photoautotrophs

## Abstract

Biological processes in the Proterozoic Ocean are often inferred from modern oxygen-deficient environments (MODEs) or from stable isotopes in preserved sediment. To-date, few MODE studies have simultaneously quantified carbon fixation genes and attendant stable isotopic signatures. Consequently, how carbon isotope patterns reflect these pathways has not been thoroughly vetted. Addressing this, we profiled planktonic productivity and quantified carbon fixation pathway genes and associated carbon isotope values of size-fractionated (0.2 – 2.7 and > 2.7 μm) particulate organic carbon values (8^13^C_POC_) from meromictic Fayetteville Green Lake, NY, USA. The high-O_2_ Calvin-Benson-Bassham (CBB) gene (*cbbL)* was most abundant in the <2.7 μm size fraction in shallow oxic and deep hypoxic waters, corresponding with cyanobacterial populations. The low-O_2_ CBB gene (*cbbM)* was most abundant near the lower oxycline boundary in the larger size fraction, coincident with purple sulfur bacteria populations. The reverse citric acid cycle gene (*aclB)* was equally abundant in both size fractions in the deepest photic zone, coinciding with green sulfur bacteria populations. Methane coenzyme reductase A (*mcrA*), of anaerobic methane cyclers, was most abundant at the lower oxycline boundary in both size fractions, coinciding with *Methanoregula* populations. 8^13^C_POC_ values overlapped with the high-O_2_ CBB fixation range except for two negative excursions near the lower oxycline boundary, likely reflecting assimilation of isotopically-depleted groundwater-derived inorganic carbon by autotrophs and acetate oxidation by sulfate-reducers. Throughout aphotic waters, 8^13^C_POC_ values of the large size fraction became 13C-enriched, likely reflecting abundant purple sulfur bacterial aggregates. Microalgal-like isotopic signatures corresponded with increases in *cbbL*, *cbbM* and *aclB*, and enrichment of exopolymer-rich prokaryotic photoautotrophs aggregates. Results suggest that 8^13^C_POC_ values of preserved sediments from areas of the Proterozoic Ocean with sulfidic photic zones may reflect a mixture of alternate carbon-fixing populations exported from the deep photic zone, challenging the paradigm that sedimentary stable carbon isotope values predominantly reflect oxygenic photosynthesis from surface waters.

## II-Introduction

Carbon stable isotope values of organic carbon (δ^13^C_OC_) preserved in ancient sediments are commonly used to infer biogeochemical carbon pathways dominating ancient water bodies (Hayes et al., 1999, Des Marais, 1997, Anbar and Knoll, 2002, Thomazo et al., 2009). They are particularly informative in sediment horizons that span extreme climatic changes, including normoxia and anoxia transitions (i.e., Ocean Anoxic Events). Use of δ^13^C_OC_ values assumes that rock-bound organic carbon is biogenic, and that it is set by: (1) the aggregate δ^13^C values of assimilated dissolved inorganic and organic carbon, (2) isotopic fractionation during fixation into eukaryote algal/prokaryotic biomass, and (3) further biological processing and isotopic fractionation as sinking POC is respired enroute to the sediments or after deposition (Hayes 1993, 2001, Werne and Hollander, 2004). Of these factors, δ^13^C_OC_ values are most influenced by autotrophic carbon-fixing enzymes that kinetically fractionate dissolved inorganic carbon (DIC) as it is converted into biomass (Emerson and Hedges, 2008).

Changes in biogeochemical carbon cycling over geologic history resulting from varying redox conditions are recorded in δ^13^C_OC_ values because key enzymes in both eukaryote algal and prokaryotic carbon fixation pathways have unique oxygen sensitivities, require different redox-sensitive metals for activation, and fractionate carbon isotopes to varying extents (Hügler and Sievert, 2010; Berg, 2011, **Table 1**). The most common application of this proxy is to search for two co-occurring trends of δ^13^C_carobonate_ values (representing ancient dissolved inorganic carbon) and δ^13^C_OC_ values of operationally-defined pools of sedimentary organic carbon (e.g., acid-insoluble organic carbon or acid-and-solvent-insoluble kerogen) over time, which approximately represents the bulk fractionation factor (ɛ). Strong negative δ^13^C_OC_ value excursions without a corresponding anomaly in δ^13^C values of carbonate indicates a shift from normoxic conditions, during which the primary carbon-fractionating process is oxygenic photoautotrophy by phytoplankton (cyanobacteria and eukaryote algae) in the euphotic zone, to “anomalous” strongly reducing conditions, during which the primary carbon-fractionating processes are non-photoautotrophic prokaryotic methane cycling and sulfate reduction. This may or may not be caused by greater organic matter burial rates. Positive excursions in both δ^13^C_OC_ and δ^13^C_carbonate_ indicates increased organic matter burial rates, which causes a drawdown of atmospheric *p*CO_2_ and more ^13^C-enriched organic matter, without strong sulfate reduction and/or methane cycling (Kump and Arthur, 1999, (Eigenbrode and Freeman, 2006).

**Table 1:**
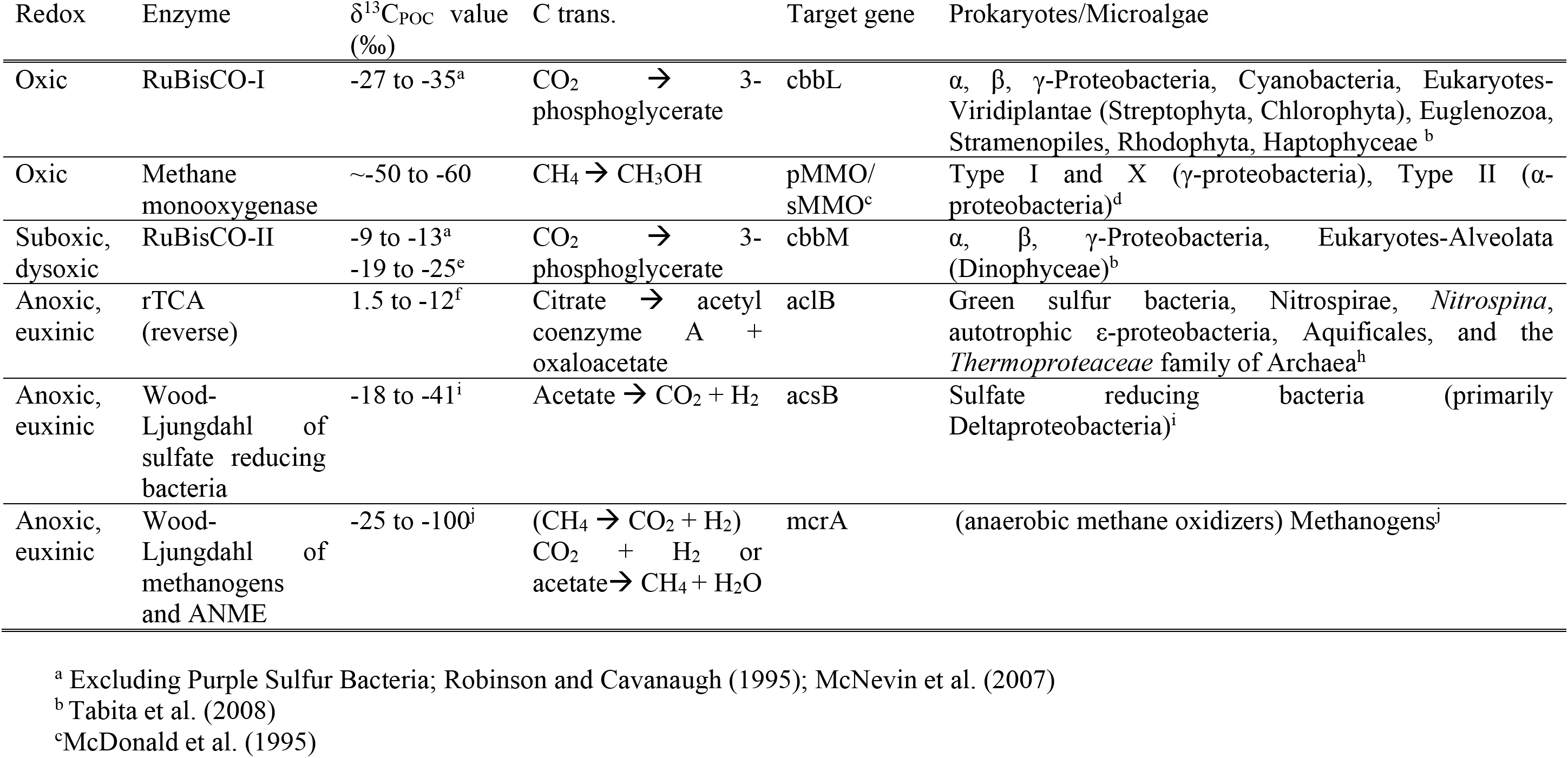
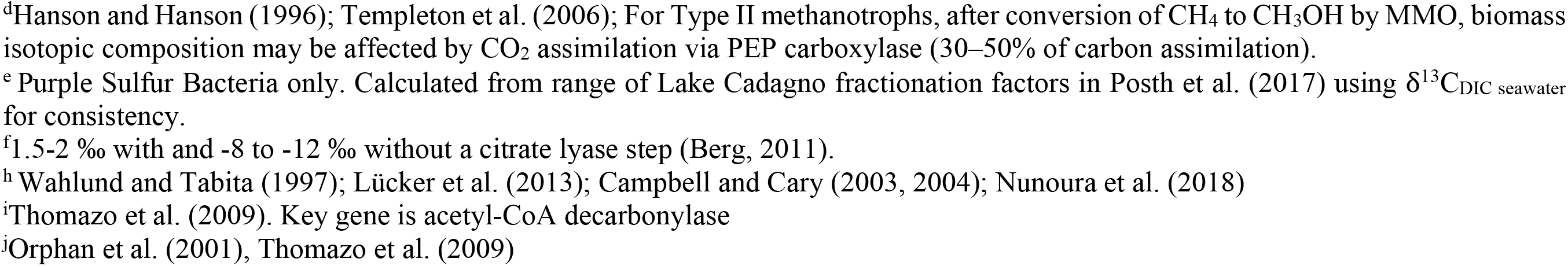
Carbon fixation pathways’ δ^13^C_POC_ values (‰), key enzymes, carbon transformations, diagnostic marker genes, and occurrence in algae and prokaryotes organized by redox condition. These values are derived from marine systems, and thus reflect fractionation ranges from dissolved inorganic carbon (DIC) in seawater: average δ^13^C_DIC seawater_ values = 1.7‰ (Cheng et al., 2019)

| Redox | Enzyme | $\delta^{13}\text{C}_{\text{POC}}$ value (‰) | C trans. | Target gene | Prokaryotes/Microalgae |
| --- | --- | --- | --- | --- | --- |
| Oxic | RuBisCO-I | -27 to -35<sup>a</sup> | $\text{CO}_2 \rightarrow$ 3-phosphoglycerate | cbbL | $\alpha$, $\beta$, γ-Proteobacteria, Cyanobacteria, Eukaryotes-Viridiplantae (Streptophyta, Chlorophyta), Euglenozoa, Stramenopiles, Rhodophyta, Haptophyceae<sup>b</sup> |
| Oxic | Methane monooxygenase | ~-50 to -60 | $\text{CH}_4 \rightarrow \text{CH}_3\text{OH}$ | pMMO/sMMO<sup>c</sup> | Type I and X (γ-proteobacteria), Type II (α-proteobacteria)<sup>d</sup> |
| Suboxic, dysoxic | RuBisCO-II | -9 to -13<sup>a</sup><br>-19 to -25<sup>e</sup> | $\text{CO}_2 \rightarrow$ 3-phosphoglycerate | cbbM | $\alpha$, $\beta$, γ-Proteobacteria, Eukaryotes-Alveolata (Dinophyceae)<sup>b</sup> |
| Anoxic, euxinic | rTCA (reverse) | 1.5 to -12<sup>f</sup> | Citrate → acetyl coenzyme A + oxaloacetate | acIB | Green sulfur bacteria, Nitrospirae, *Nitrospina*, autotrophic ε-proteobacteria, Aquificales, and the *Thermoproteaceae* family of Archaea<sup>h</sup> |
| Anoxic, euxinic | Wood-Ljungdahl of sulfate reducing bacteria | -18 to -41<sup>i</sup> | Acetate $\rightarrow \text{CO}_2 + \text{H}_2$ | acsB | Sulfate reducing bacteria (primarily Deltaproteobacteria)<sup>i</sup> |
| Anoxic, euxinic | Wood-Ljungdahl of methanogens and ANME | -25 to -100<sup>j</sup> | $(\text{CH}_4 \rightarrow \text{CO}_2 + \text{H}_2)$<br>$\text{CO}_2 + \text{H}_2$ or<br>acetate $\rightarrow \text{CH}_4 + \text{H}_2\text{O}$ | mcrA | (anaerobic methane oxidizers) Methanogens<sup>j</sup> |

<sup>a</sup> Excluding Purple Sulfur Bacteria; Robinson and Cavanaugh (1995); McNevin et al. (2007) <sup>b</sup> Tabita et al. (2008) <sup>c</sup> McDonald et al. (1995)
<sup>d</sup>Hanson and Hanson (1996); Templeton et al. (2006); For Type II methanotrophs, after conversion of CH<sub>4</sub> to CH<sub>3</sub>OH by MMO, biomass isotopic composition may be affected by CO<sub>2</sub> assimilation via PEP carboxylase (30–50% of carbon assimilation). <sup>e</sup> Purple Sulfur Bacteria only. Calculated from range of Lake Cadagno fractionation factors in Posth et al. (2017) using $\delta^{13}\text{C}_{\text{DIC}}$ seawater for consistency. <sup>f</sup>1.5-2 ‰ with and -8 to -12 ‰ without a citrate lyase step (Berg, 2011). <sup>h</sup> Wahlund and Tabita (1997); Lückner et al. (2013); Campbell and Cary (2003, 2004); Nunoura et al. (2018) <sup>i</sup>Thomazo et al. (2009). Key gene is acetyl-CoA decarbonylase <sup>j</sup>Orphan et al. (2001), Thomazo et al. (2009)

Although it has been suggested that a bulk fractionation factor greater than 32‰ indicates high contributions by anaerobic photo- and/or chemo-autotrophs (Hayes et al., 1999) to the preserved organic carbon pool, the 28-32‰ indicative of maximal contributions from oxygenic photoautotrophs are difficult to distinguish from values at the low end of the greater-than-32‰ range, considering the analytical error of δ^13^C mass spectrometry measurements. So, in practice, ɛ values in this potentially overlapping range that may be indicative of anaerobic autotrophs are typically interpreted as representing oxygenic photosynthesis. As a result, prior studies may have overlooked the existence of ancient oxygen-deficient water bodies that hosted stratified populations of photo- and chemoautotrophs, each with distinctive carbon fixation pathways that produce characteristic stable isotope patterns as observed in modern oxygen-deficient aquatic environments (MODEs) (e.g., Hügler and Sievert, 2010, Ruiz-Fernandez et al., 2020) (**Table 1**).

In shallow oxygenated ocean water, form I of the ribulose-1,5-bisphosphate carboxylase/oxygenase (RuBisCO) enzyme characteristic of the high-O_2_ Calvin-Benson-Bassham (CBB) cycle, and present in all microalgae and cyanobacteria, likely dominates, producing POC with δ^13^C values that vary between -35 and -27‰ from marine DIC (δ^13^C_DIC_ values∼1.7‰; Cheng et al., 2019)(Tabita et al., 2008). Within oxyclines of stratified water bodies, where conditions range from oxic to suboxic, δ^13^C values may reflect a mixture of high-O_2_ RuBisCO form I and the low-O_2_ RuBisCO form II, found in photo-(δ^13^C_POC_ values -25 to -19‰;) and chemo-autotrophic (δ^13^C_POC_ values = -13 to -9‰), sulfur-oxidizing gammaproteobacteria _(_Canfield et al., 2010, Imhoff, 1995, Posth et al., 2017). While O_2_-sensitive, the low-O_2_ CBB pathway includes a protective enzyme that reduces O_2_ to H_2_O (Probst et al., 2017). Thus, microbes fixing carbon through the low-O_2_ CBB pathway can thrive in the presence of micromolar oxygen concentrations and/or transient oxygenation events. Prokaryotes fixing carbon through the reverse citric acid cycle (rTCA) using the ATP citrate lyase enzyme (*aclB*) do not have such an oxygen protection mechanism. Therefore, these microbes, including the sulfur-oxidizing chemoautotrophic epsilonproteobacteria and photoautotrophic green sulfur bacteria, reside below the oxycline in deeper, quiescent sulfidic waters (Lin et al., 2006, Grote et al., 2008, Imhoff, 1995). Finally, within a narrow range of redox and chemical conditions, the strongly carbon-fractionating methane-cycling prokaryotes may have the greatest impact on δ^13^C values. Methyl coenzyme reductase A in the reductive acetyl coenzyme A (Wood-Ljungdahl) pathway is utilized by methanogenic Euryarchaeota in the forward direction and in the reverse direction by anaerobic methane oxidizing Archaea (ANMEs) (Hallam et al., 2004). Enzymes involved in methane cycling are hyper-sensitive to oxygen and depend on bioavailability of redox-sensitive metals (e.g., Mo, Co, Ni, Fe) that are insoluble under oxygenated conditions or when exposed to moderate to high concentrations of hydrogen sulfide (Berg, 2011, Momper et al., 2017). These limitations restrict ANME and methanogen distributions to the sulfate-methane transition zone in sediments or just below oxycline boundary in anoxic water columns, where conditions are reducing but hydrogen sulfide is either scarce or absent (Jorgensen et al., 2001, Dhillon et al., 2005).

It is axiomatic that relationships observed in modern environments offer a lens into the geologic past and enable a better understanding of Earth’s history. The possibility that δ^13^C_OC_ values within the range diagnostic of oxygenic photosynthesis in ancient sediment integrates biomass carbon produced through the many carbon fixation pathways supported in oxygen-deficient water bodies has received attention predominantly from an organic geochemical perspective (Johnston et al., 2009). Models groundtruthed with compound-specific isotope studies have revealed this caveat in paleobiogeochemical studies of past water columns (Fulton et al., 2018).

A cross-disciplinary exploration of relationships between microalgal and prokaryotic carbon fixation pathways and isotope ratios of potentially sedimenting materials can expand our understanding of complex biogeochemical carbon cycling on ancient Earth. However, studies that combine spatial quantitation of carbon fixation genes and measurements of δ^13^C_POC_ values in MODEs are rare. We hypothesize that organic material preserved in Proterozoic sediments, deposited in waters with a sulfidic photic zone, may reflect an admixture of carbon fixation pathways while superficially resembling the cyanobacterial δ^13^C signatures. To bridge this knowledge gap and test our hypothesis, we profiled carbon fixation pathway marker genes (**Table ^1^**) by quantitative polymerase chain reactions and measured δ^13^C_POC_ through all redox zones of Fayetteville Green Lake (FGL), one of the world’s most extensively studied meromictic lakes. Because cells that aggregate or associate with particles are more commonly buried in sediments owing to their faster settling velocities relative to small suspended individual cells (Alldredge and Gotschalk, 1988), we analyzed size-fractionated particulate matter. We compare current results with size-fractionated autotrophic and methane-cycling phylotype distributions previously determined by 16S rRNA amplicon sequencing (Cohen et al., 2023). To provide a biogeochemical context, we profiled hydrographic and chemical features and microbial production rates.

Fayetteville Green Lake, NY, USA (FGL hereafter) is 52 m deep, sulfidic (euxinic), meromictic, and located in the Oswego River-Lake Ontario watershed. FGL has been studied extensively, especially to gain insight into dominant biogeochemical cycling pathways in the ancient ocean during periods of widespread and prolonged anoxia (e.g., Zerkle et al., 2010, Havig et al., 2018, Fulton et al., 2018). Although a freshwater lake, FGL has high sulfate concentrations because groundwater intrusions pass through gypsum-bearing sedimentary rock (Brunskill and Ludlam, 1969, Torgerson et al., 1981). Consequently, FGL’s vertical biogeochemical zones are not unlike marine anoxic basins (e.g., the Cariaco Basin and the Black Sea). These include an oxygenated mixed layer (mixolimnion) and a sulfidic deep layer (monimolimnion) separated by an oxycline with redox conditions spanning oxic, hypoxic, and suboxic (Zerkle et al., 2010, Havig et al., 2015, Cohen et al., 2023). Because the photic zone extends from oxic to sulfidic waters, it contains stratified photoautotrophic populations that employ a range of carbon fixation pathways (**Table 1**). The shallow oxic and deep hypoxic zones are inhabited by cyanobacterial populations that induce calcium carbonate precipitation (so-called “whiting events”; Thompson et al., 1990).

The lower photic zone is sulfidic, has elevated microbial activity, biomass, and turbidity, and is dominated by purple and green anoxygenic sulfur-oxidizing photoautotrophs (Cohen et al, 2023). The aphotic monimolimnion is euxinic and methanic. The extremely light 8^13^C_CH4_ values (∼-100 ‰) indicate that the CH_4_ is biogenic and that sediments are inhabited by strongly carbon-fractionating methane-cycling microorganisms (**Table 1**, Havig et al., 2018). The structure of FGL’s photic zone and underlying euxinic and methanic aphotic zone is thought to be more like large expanses of the ancient ocean than modern marine anoxic basins (Havig et al., 2018).

## III-Materials and Methods

### Field sampling

For this study, the deepest part of the FGL water column (43°03’01.9“N, 75°57’58.9”W) was sampled on 17-21 July 2017 and 28 July-4 August 2018. Sampling depths spanned all redox zones (10 – 40 m), with finer vertical resolution near the lower oxycline boundary, where vertical biogeochemical gradients are steep. To determine the physico-chemical structure, we profiled dissolved oxygen, light scattering (turbidity), fluorometric phycoerythrin and chlorophyll-a concentrations, and salinity using a YSI EXO1 sensor package. Total microbial cell and hydrogen sulfide concentrations were measured in discrete samples as described in Cohen et al., 2023.

During July 2017, total inorganic carbon assimilation and bacterial heterotrophic production (BHP) were profiled throughout the entire water column using radioactive tracers (^14^C-bicarbonate and ^3^H-leucine, respectively; Cohen et al., 2023). During July 2018, inorganic carbon assimilation was profiled from 19-25 m at a finer vertical resolution and proportions of dark (chemoautotrophic) and light (photoautotrophic) assimilation were determined. Detailed sample retrieval and processing protocols are presented in Cohen et al. (2023). Briefly, samples were retrieved by directly pumping water from the sample depth into vials after overflowing three times and incubated on site. Samples from photic depths and killed controls were incubated on floating racks in mesh bags layered to mimic in situ illumination within open water incubators to maintain in situ temperatures. Samples from aphotic depths or dedicated to dark inorganic carbon assimilation measurements were placed in opaque bags at the bottom of the incubator. Terminated incubations were stored refrigerated in the dark until processing. Sample processing, radioactivity measurements by liquid scintillation counting, and the conversion of sample radioactivity to inorganic carbon assimilation and bacterial heterotrophic production rates were performed as previously described (Taylor et al. 2001, Cohen et al., 2023)

Discrete samples were collected and separated into particle-associated (PA) and free-living (FL) size fractions by in-line filtration through >2.7 µm GF/F and 0.2 µm polycarbonate Sterivex™ filters for DNA recovery and analyses during July 2017, as described elsewhere (Cohen et al., 2023). It should be recognized that the PA fraction may also include cell aggregates, prokaryotes symbiotically associated with larger protists and zooplankton, and detrital terrestrial plant matter. By the same token, the FL fraction may include some particle-associated prokaryotes washed free of their particles during sample processing. Samples for stable carbon isotope measurements collected during July 2018 correspond to July 2017 DNA sample depths. PA and FL size fractions were obtained by filtering samples from each depth sequentially through pre-combusted 2.7 µm and 0.7 µm Whatman GF/F flat filters using a peristaltic pump. We acknowledge that the nominal pore size of the FL GF/F filter is greater than the Sterivex™ filter used to collect the DNA samples, but it has been shown that these filter-types retain approximately the same concentration of environmental DNA (Minamoto et al., 2015). Filters were flash-frozen with dry ice and stored at -20°C until processing.

### Carbon fixation gene quantification

Quantitative polymerase chain reaction (qPCR) assays were optimized using pooled (equal volumes of all collected samples) environmental DNA that was extracted, aliquoted, and stored at -80°C in 2017 (Cohen et al., 2023) as the template. All qPCR was performed in 25 µL reactions using Lucigen FailSafe™ reagents except for the *mcrA* gene, for which 20 µL reactions were prepared using Applied Biosystems™ PowerUp™ SYBR™ Green Master Mix (ThermoFisher Scientific^TM^). One µL of template was used with both protocols. Because sample DNA concentrations varied widely among samples (from 10 ng/ µL in the mixolimnion to 100s of ng/ µL near the lower oxycline boundary), dilutions required per sample were empirically determined during inhibition tests. All qPCR was performed with an Applied Biosystems™ QuantStudio 6 Real Time PCR machine (ThermoFisher Scientific^TM^) using ROX as the passive reference dye and SYBR-Green I as the reporter dye. All non-quantitative PCR was performed using a Labnet MultiGene™ Optimax thermal cycler.

For each assay, PCR was first performed to optimize reaction chemistry and annealing temperature according to each of the manufacturer’s instructions, using the published primer concentrations, and always including a non-template control (NTC). Resulting PCR products and a reference ladder were qualitatively assessed after running gel electrophoresis on either 1% agarose gels in 1x TAE buffer (product size >250 base pairs) or 2% agarose gels in 1x TBE buffer (product size < 250 base pairs).

Quantitative standards were created from PCR products of pooled environmental DNA using the optimized thermal profiles. If gel visualization revealed poor PCR product quality, amplicons were first purified using a Zymo Research Genomic DNA Clean & Concentrator™ kit. To create standard stocks of a desired concentration (copies/μL), the NTC-corrected dsDNA content of the pooled DNA PCR product was quantified by fluorescence using a Quant-iT™ PicoGreen™ dsDNA assay kit (Invitrogen™) as in Blotta et al. (2005). The NTC-corrected dsDNA content of the PCR product was converted to gene copies/µL using the published number of base pairs of the PCR product, the average molar mass per base pair of dsDNA (660 g/mol), and Avogadro’s number (6.03 x 10^23^ molecules/mol) as conversion factors. Stocks were created by diluting product with 1x TE. Aliquots were stored at -80°C, with each aliquot subject to no more than 6 freeze-thaw cycles.

qPCR primer concentrations for PowerUp™ reagent assays and primer-SYBR Green I concentration combinations for FailSafe™ assays were optimized using 5-7 point standard curves until the R^2^ of the best-fit line to ΔR_n_ versus log_10_ concentration were >0.98 and reaction efficiencies were between 90 and 110%. Product melting temperatures, primer dimers, and non-specific amplification were evaluated from melt curves. Finalized reaction chemistries and thermal profiles are shown in **Table 2**. Using optimized reaction chemistries, potential for PCR inhibition was determined by running reactions of serially diluted environmental DNA samples. Appropriately-diluted environmental samples, including full field and laboratory procedural blanks, were run in triplicate with a full standard curve. Diagnostic PCRs were prepared for each environmental sample to cross-check for the presence of the gene.

**Table 2:**
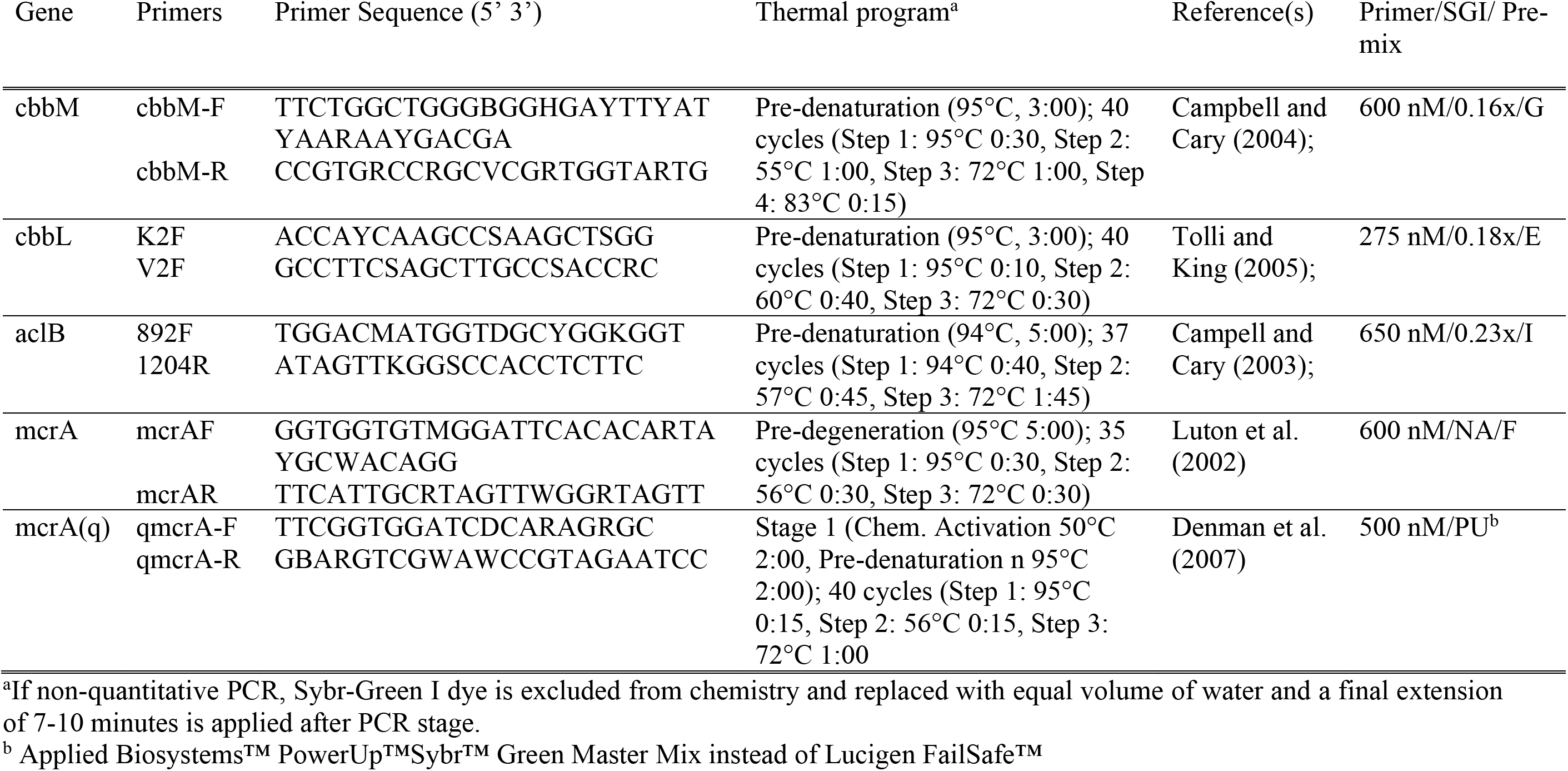
Carbon fixation pathway marker genes targeted by qPCR, primer information, and finalized reaction chemistry.

### Carbon stable isotope measurements

After lyophilization, subsamples were punched out from filters. Filter subsamples were placed in silver capsules and treated by sequential HCl fumigation in a desiccator to remove residual inorganic carbon. The first fumigation was performed using 20% HCl for 24 hours after rewetting subsamples with 25 µL of 1N HCl. The second fumigation was performed using 37% HCl for 24 hours after rewetting subsamples with 25 µL ddH_2_O. These subsamples were stored in a 60°C drying oven until elemental analyzer-isotope ratio mass spectrometry (EA-IRMS) analysis.

All EA-IRMS measurements were made in duplicate and bracketed every 5 samples with standards on a ThermoScientific^TM^ Delta V Plus IRMS coupled to an EA Isolink elemental analyzer and Costech Zero-blank autosampler in the Department of Geosciences at Stony Brook University. Every EA-IRMS run also included true (sampling) blanks. Stable carbon isotope ratios are reported as per mil (‰) in delta notation:

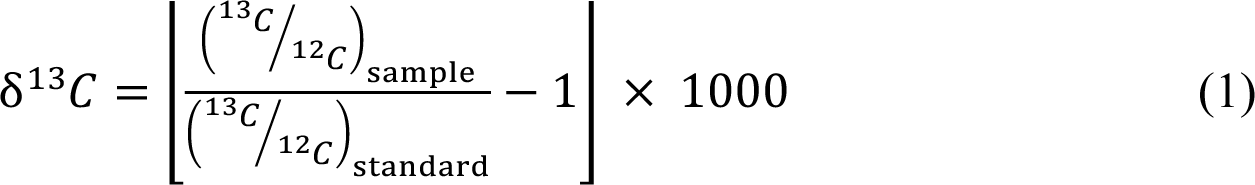

where the standard is the Vienna Pee Dee Belemnite. Standard deviations of δ^13^C measurements (n=5) of the bracketing certified standards USGS65, IU-L glutamic acid, and IAEA-600 were ±0.04, ±0.03, ±0.03 ‰, respectively. In our natural samples, average standard deviations for measurements of true δ^13^C_POC_ replicates of the PA and FL fractions were 0.50‰ and 0.62‰ for the PA and FL fractions, respectively.

## IV-Results

### Water column structure

We profiled the physico-chemical structure and major microbiological features of FGL during July 2017 and July 2018 to provide a context for our qPCR and δ^13^C profiles. Observations from July 2018 were not significantly different from those of July 2017 (**Figs. 1-3**). During both field campaigns, the mixolimnion had a shallow (∼3 m) mixed layer comprised mostly of surficial run-off water, indicated by a uniform dissolved oxygen concentration and low salinity (**Figs. 1,2**). Immediately below 3 m, dissolved oxygen concentrations rose to a maximum of 400 μM (supersaturated) due to oxygenic photosynthesis by shallow cyanobacteria and diatom populations, then declined to ∼300 μM at the upper oxycline boundary (15 m). The lower oxycline boundary, defined by the first appearance of H_2_S, was located at 20 m. H_2_S concentrations during both years steadily increased with depth in the monimolimnion to ∼2 mM at 40 m (**Fig. 1**).

**Figure 1:**
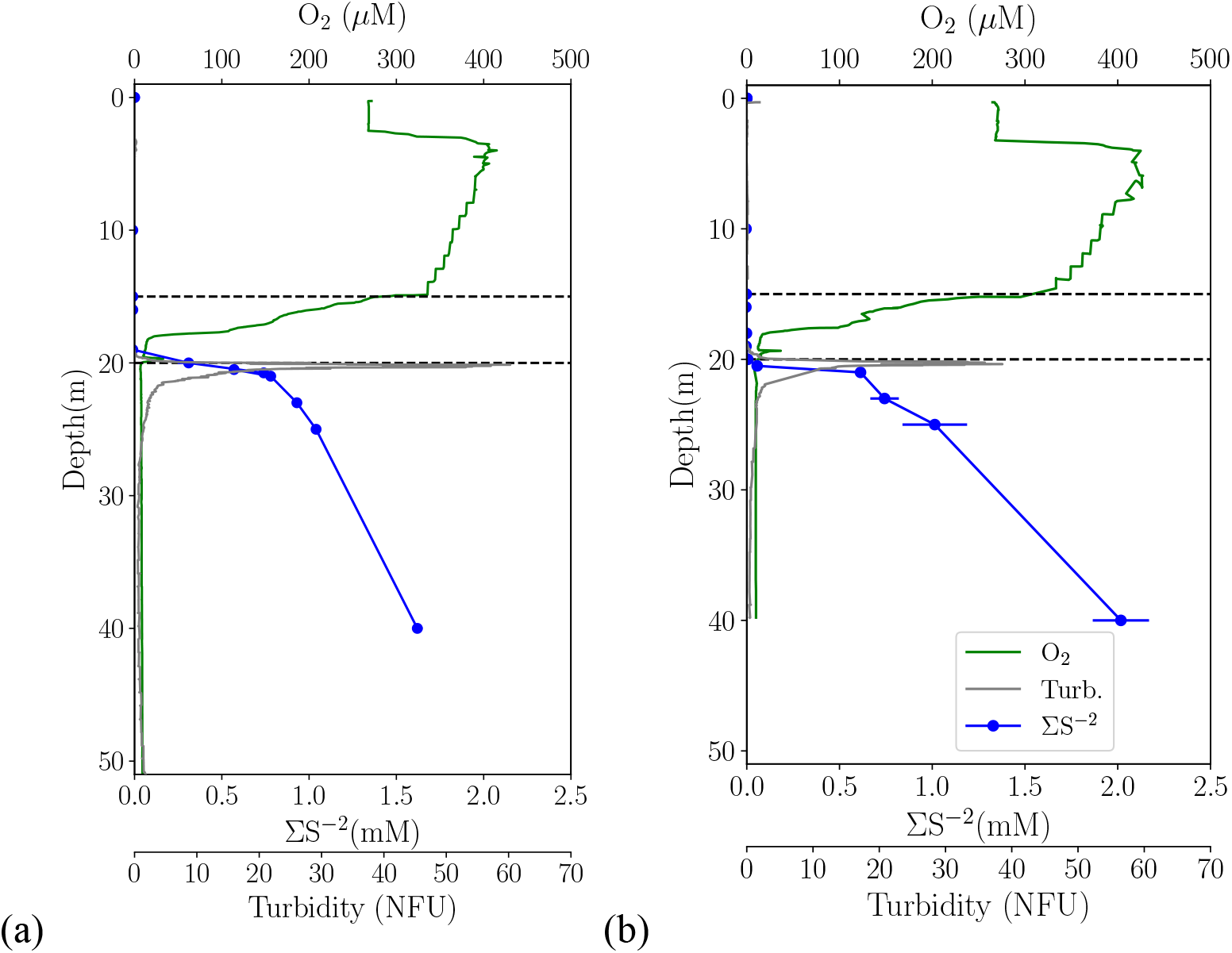
Vertical profiles of redox conditions and turbidity during (a) July 2017 and July 2018 (b). Broken lines indicate oxycline boundaries. Error bars in ∑S^-2^ profiles represent ± 1 S.D. of the mean of triplicate analyses. Error bars fall within the size of the symbols for July 2017.

**Figure 2:**
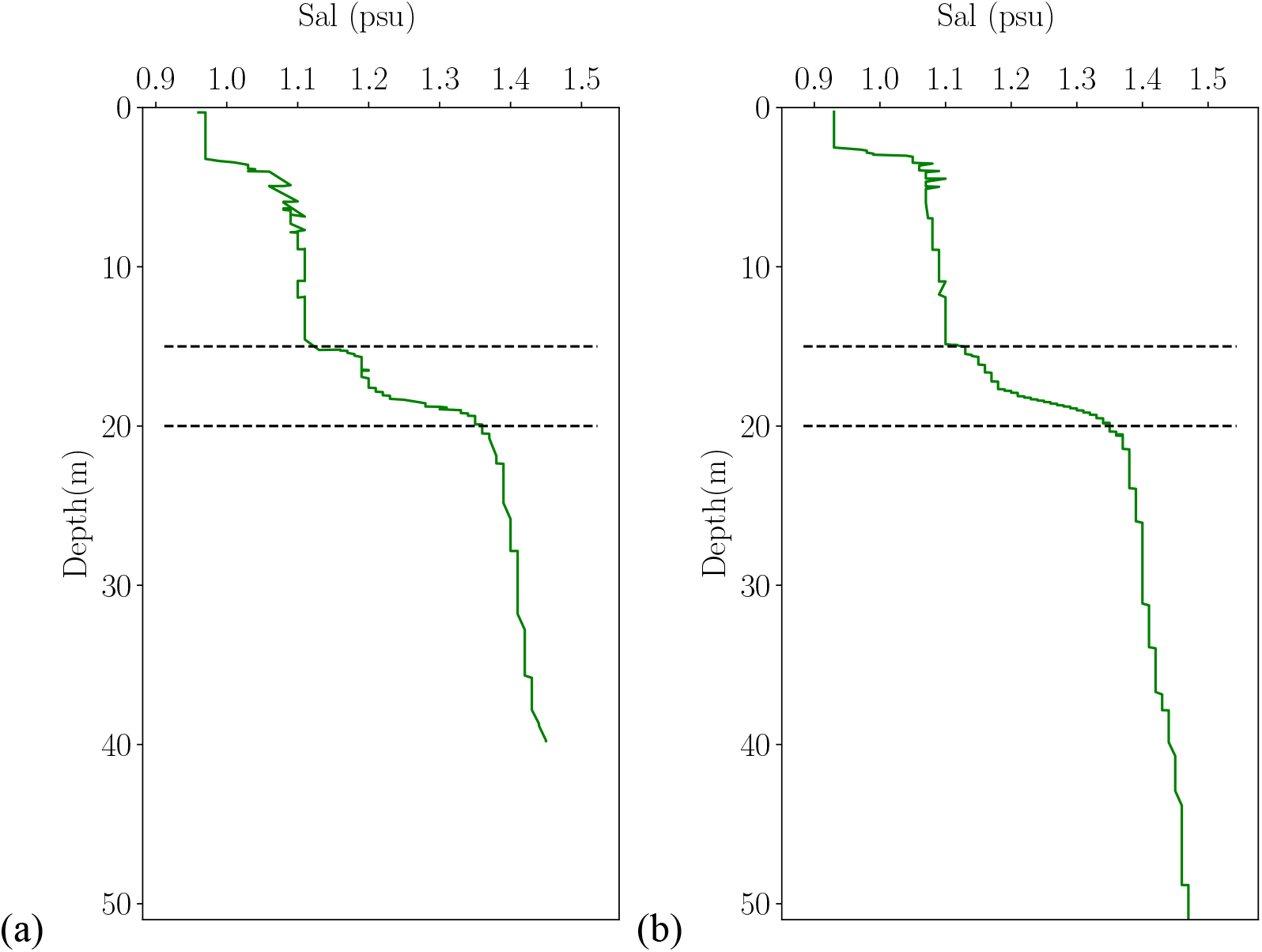
Vertical profiles of salinity measured during (a) July 2017 and July 2018 (b) using a YSI EXO1 sensor package. Broken lines indicate oxycline boundaries.

**Figure 3:**
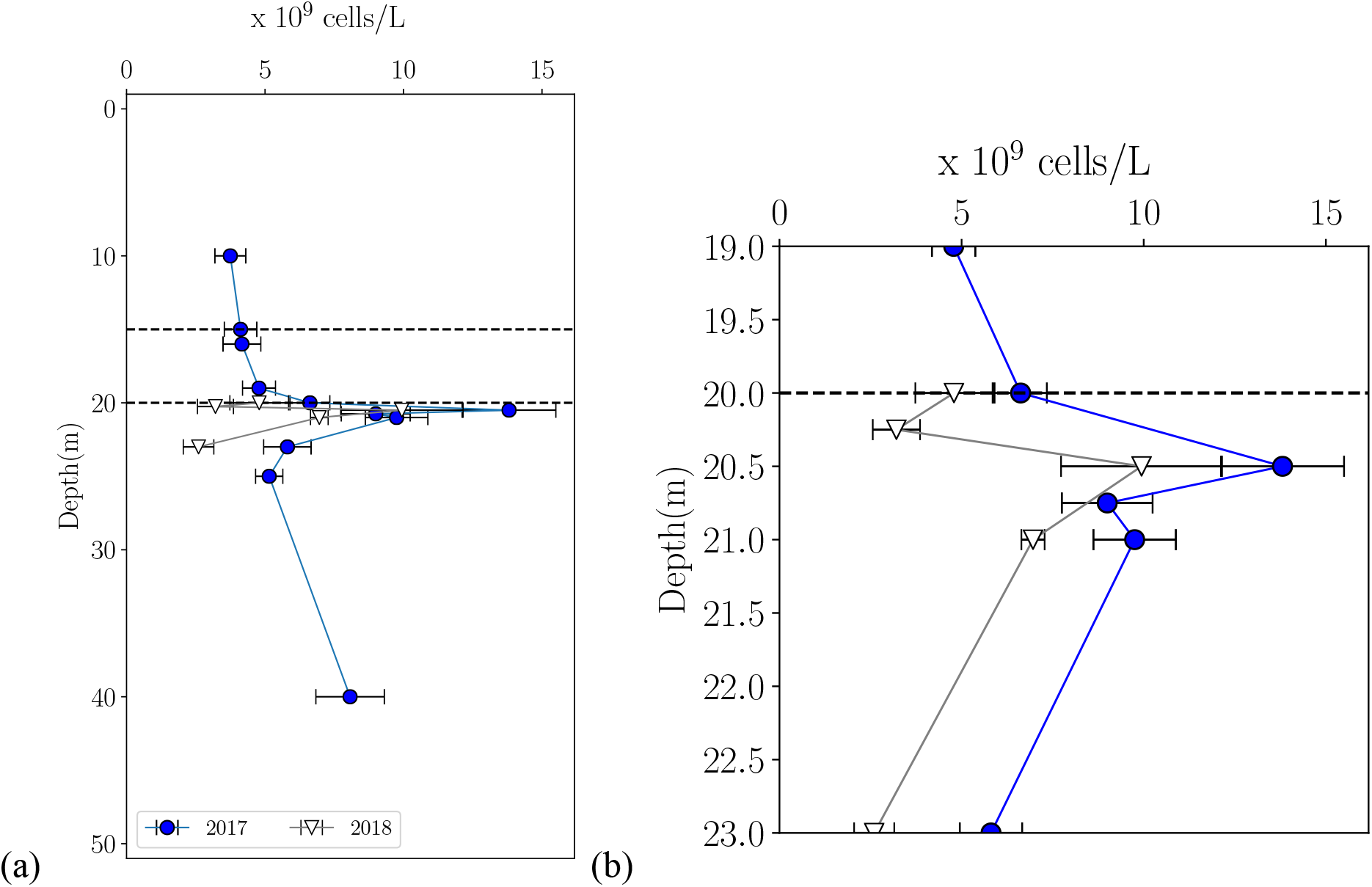
Vertical profiles of DAPI-stainable total prokaryotic cell concentrations during July 2017 (full profile) (a) and in July 2018 (profiled across lower oxycline and upper monimolimnion) (b). Broken lines indicate oxycline boundaries. Error bars represent ± 1 S.D. of duplicates.

A very narrow light-scattering layer, recognized by high measured turbidity (total particles), was detected between 19 and 23 m and coincides with elevated prokaryotic cell concentrations and total inorganic carbon assimilation (ICA) rates observed both years **(Figs.** **Figure 1, 3, 4).** We determined that the majority of ICA throughout this layer in 2018 was photoautotrophic based on ^14^C-bicarbonate assimilation in light and dark incubations (**Fig. 4b**). During both years, the shallowest portion of the light-scattering layer (19-19.75 m) coincided with a very narrow oxygen peak (**Fig. 1**) and maximum phycoerythrin concentrations, indicating local oxygen production from a deep cyanobacteria population (**Fig. 5**). The light-scattering maximum (20.25-20.5 m) aligned with maximum prokaryotic cell concentrations and maximum light and dark autotrophic ICA rates (**Figs. 3**, **Fig. 4**). Water in this layer was visibly purple, indicating the presence of a dense population of anoxygenic purple sulfur bacteria (**Fig. S1**). Abundant anoxygenic green sulfur bacteria were observed in bright-field micrographs of 21-23 m samples (**Fig. S2a**).

**Figure 4:**
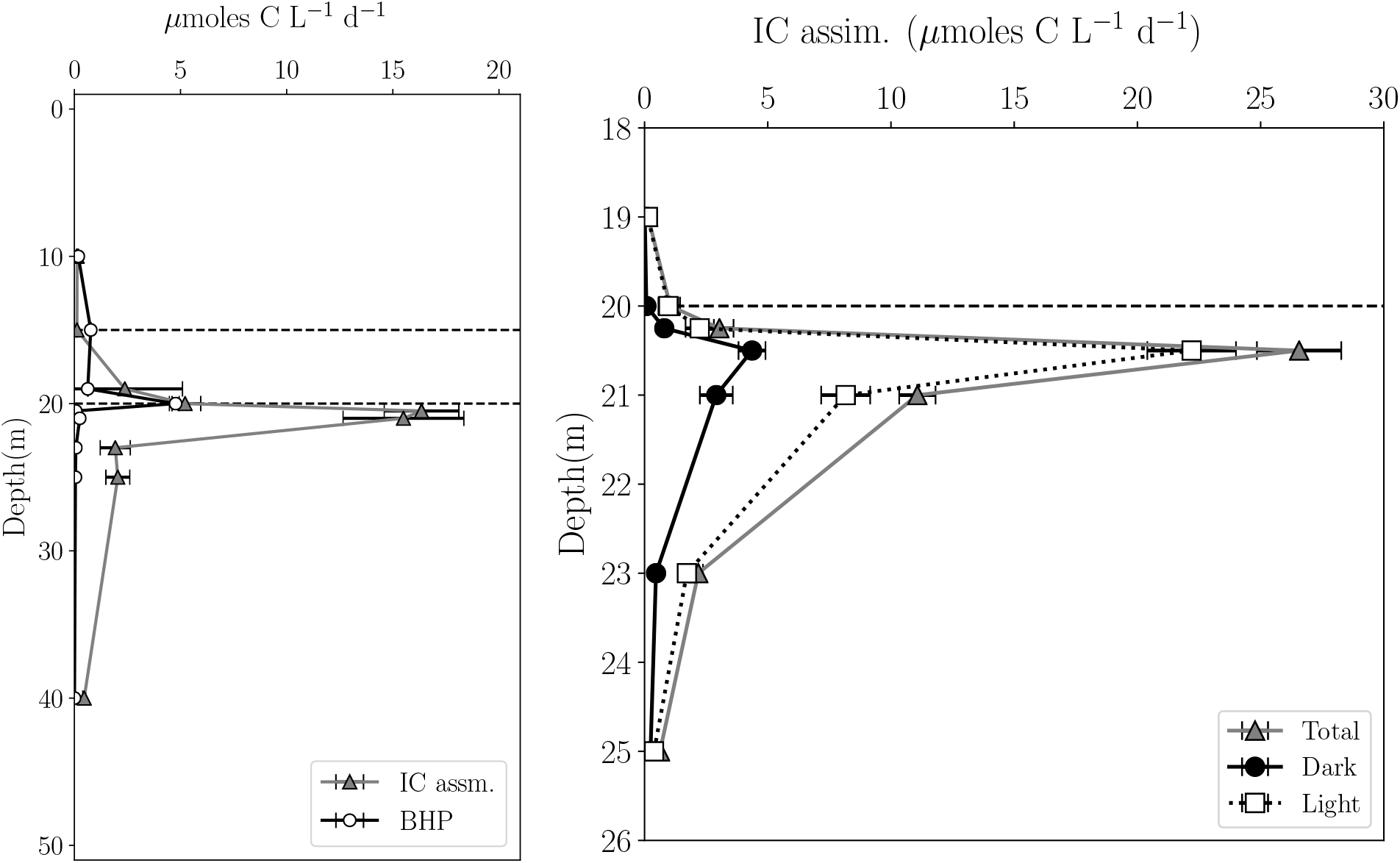
Vertical profiles of total inorganic carbon assimilation (ICA) and bacterial heterotrophic production (BHP) in July 2017 (a). Total, dark, and light ICA across the lower oxycline and upper monimolimnion at fine vertical resolution in July 2018 (b). Broken lines indicate oxycline boundaries. Error bars represent ± 1 S.D. of the mean of triplicate analyses.

**Figure 5:**
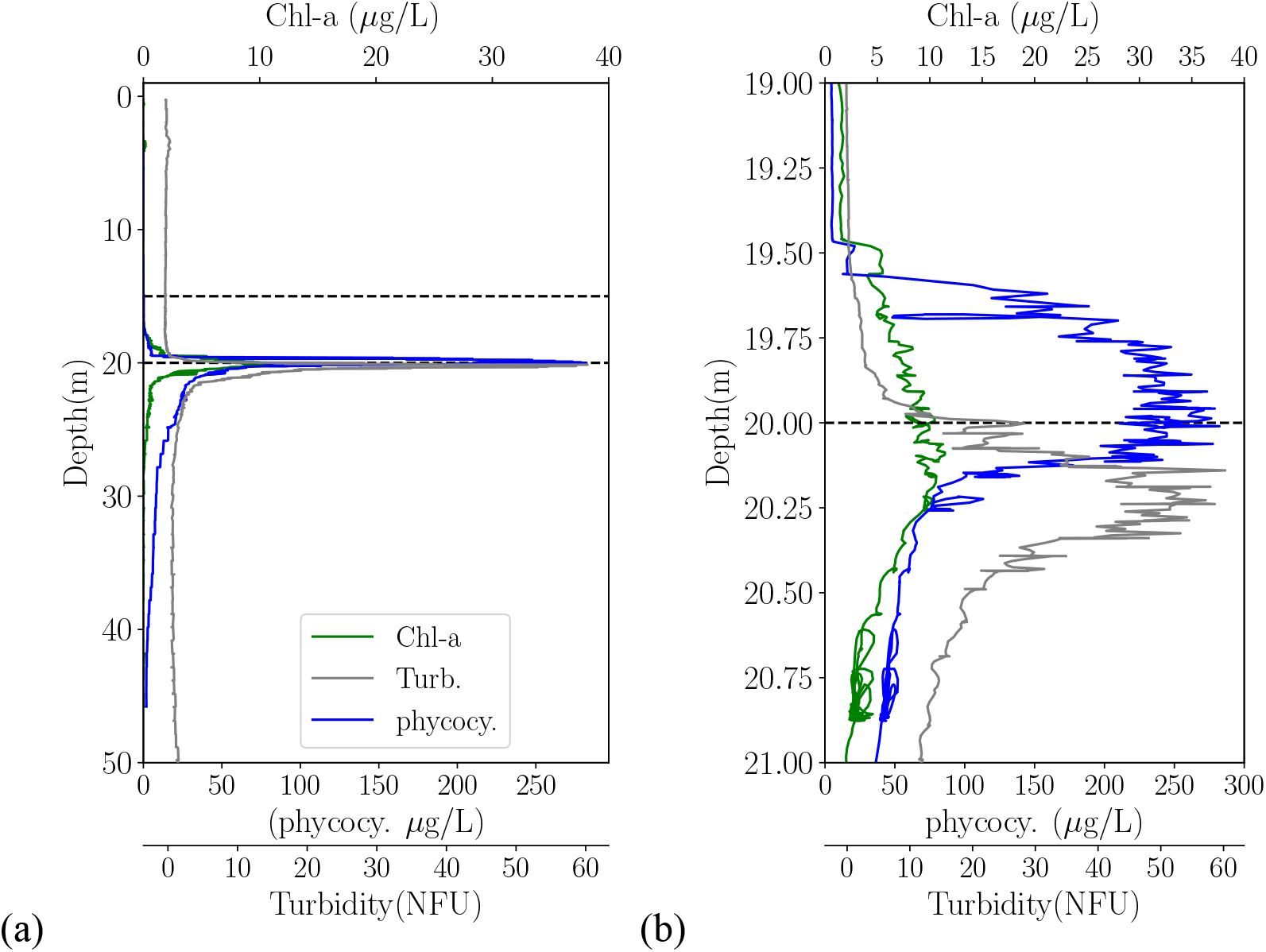
Vertical profiles of phycoerythrin fluorescence (cyanobacteria), chlorophyll-a fluorescence (all algae), and turbidity (total particles) during July 2017 of the entire water column (a) and at finer resolution across the lower oxycline (b).

### Carbon Stable Isotopes

POC in the PA size fraction was consistently more ^13^C-depleted by 2-4‰ than that recovered in the FL fraction throughout the water column, except at 20 m (lower oxycline boundary) (**Fig. 6**). From 18 to 19 m, where the deep cyanobacteria population lives, a small enrichment in ^13^C values (from -34.7‰ to -33.7‰) was observed in the FL fraction. At the lower oxycline boundary (20 m), where photoautotrophic purple and green sulfur bacteria first appear, the δ^13^C of both size fractions were most similar and were strongly ^13^C-depleted (δ^13^C_POC_ values =-41.9‰ and -39.4‰ in PA and FL fractions, respectively). At 20.5 m, where sulfur-oxidizing photoautotrophic purple sulfur bacteria and chemoautotrophic epsilonproteobacteria were previously observed to be most abundant (Cohen et al., 2021, 2023), δ^13^C_POC_ values of the two size fractions differed the most; PA fraction δ^13^C_POC_ values reached their lowest value (−42.3‰) while in the FL fraction δ^13^C_POC_ values returned to mid-oxycline values (−35.1‰). Finally, local δ^13^C_POC_ value maxima in the FL (−35.3‰) and PA (−39.8‰) fractions were evident at 21 m, where photoautotrophic green sulfur bacteria were most abundant. Below the photic zone, δ^13^C_POC_ values of both size fractions converged with depth primarily because the PA fraction was ^13^C-enriched to -37.0‰, while the FL fraction’s δ^13^C_POC_ values remained at -35.7‰. The deepest samples taken for this study were recovered from 40 m, 12 m above the lakebed.

**Figure 6:**
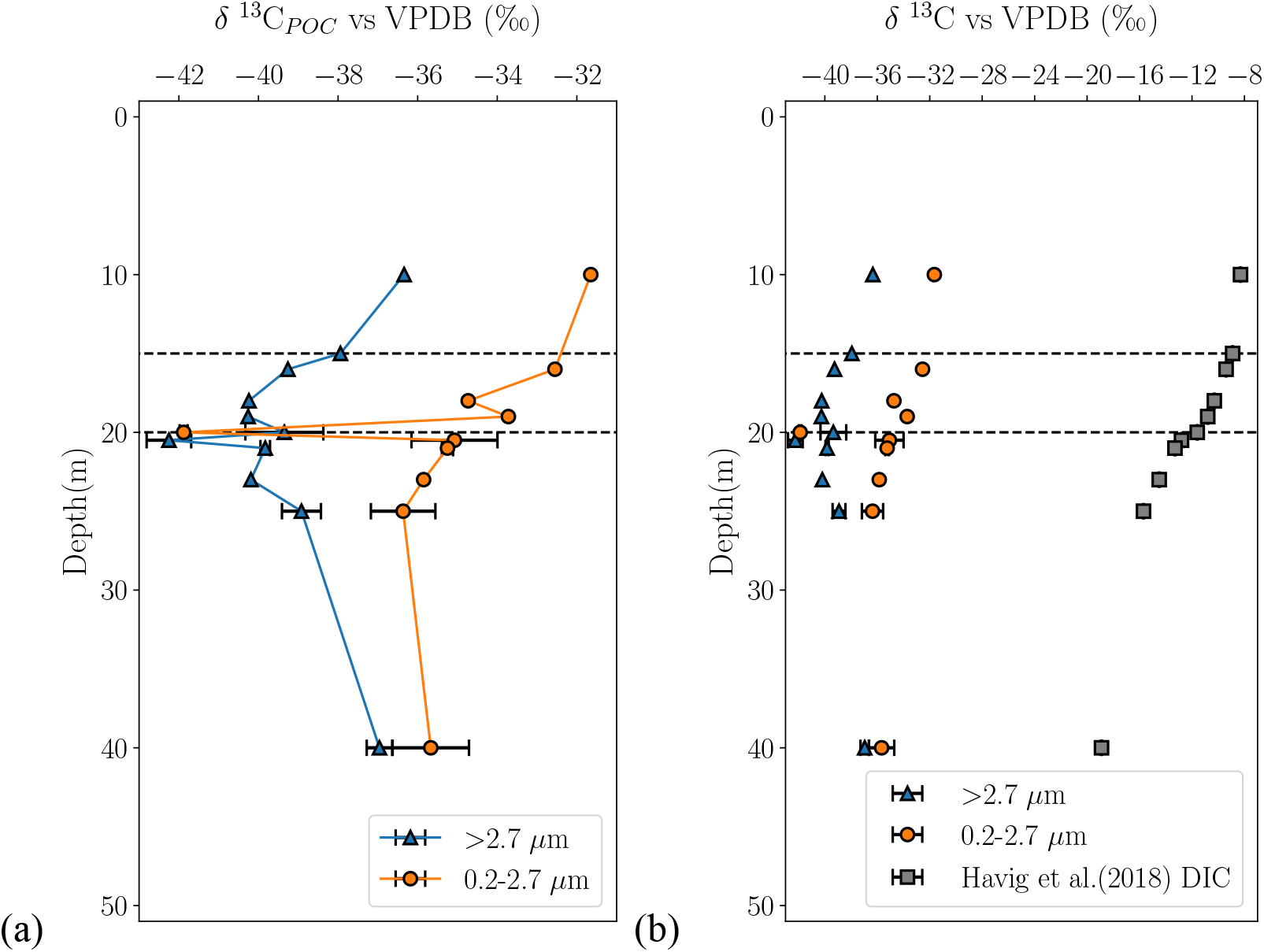
(a) Depth profile of particulate organic carbon (POC) stable isotopic composition of FL (0.2-2.7 µm) and PA (>2.7 µm) size fractions during July 2018. (b) Data presented in (a) alongside dissolved inorganic carbon (DIC) stable isotopic composition replotted from Havig et al. (2018). Broken lines indicate oxycline boundaries. Error bars represent ± 1 S.D. of duplicate analyses.

### Carbon fixation pathway genes

Size-fractionated *cbbL, cbbM*, *aclB,* and *mcrA* functional gene depth profiles reflected distributions of cyanobacterial, purple sulfur and green sulfur bacterial photoautotrophic populations, and the methanogen *Methanoregula*, respectively, determined from amplicon libraries prepared from the same samples (Cohen et al., 2023) and agreed with microscopic and sensor observations (**Fig. 7**). Copy numbers of *cbbL* in the FL fraction were more abundant than in the PA fraction at the 10 and 19.75 m maxima, the observed depths of the shallow and deep cyanobacterial populations detected by phyocerthyrin fluorescence (**Fig. 7a**). Elsewhere, copies were about equally abundant in both size fractions. Copy numbers of *cbbM* in the PA fraction were much more abundant than in the FL fraction throughout most of the water column (**Fig. 7b**), consistent with microscopic observations of purple sulfur bacteria appearing mostly in aggregates (**Fig. S2b**). Maximum *cbbM* copy numbers occurred just below the lower oxycline boundary and aligned well with turbidity maxima (**Fig. 1**), total microbial cell concentrations (**Fig. 3**), and total inorganic carbon assimilation rates (**Fig. 4**) associated with the purple sulfur bacteria population (**Fig. S1**). Inorganic carbon assimilation rates were approximately 3-fold greater than heterotrophic uptake rates (calculated as approximately 3-fold the bacterial heterotrophic production rate; REF) at the same depth, and most of the inorganic carbon assimilation is photoautotrophic, suggesting that the purple sulfur bacteria contribute the majority of the POC at this depth. Copy numbers of *aclB* were equally represented in PA and FL fractions, with maximum copy numbers aligning well with the green sulfur bacteria population in the deep photic zone (**Figs. 7c, S2a).** Therefore, chemoautotrophs’ influence on δ^13^C_POC_ is likely minimal relative to the highly abundant photoautotrophs in the photic zone. *mcrA* was equally represented in both size fractions (**Fig. 7d**). We caution that *mcrA* depth profiles should be interpreted for depth trends, but not absolute gene copy numbers. The gene was quantified with different reagents than the other genes, and standard curve and sample measurements were less consistent than those of the other quantitative methods. However, we confirmed the presence of the gene in samples by gel electrophoresis following standard PCR amplification using the same reagents as the other qPCR assays (**Table S1**).

**Figure 7:**
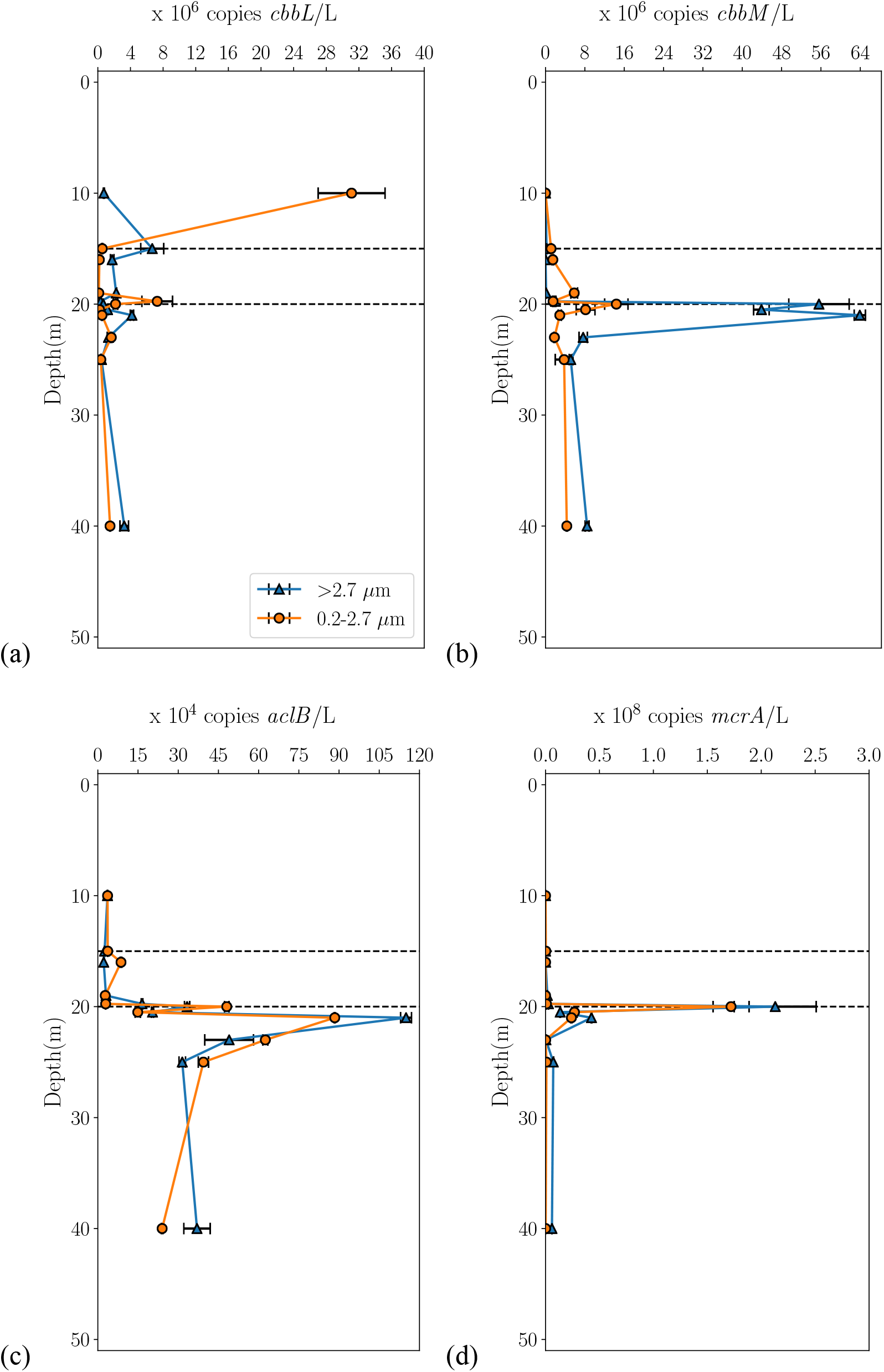
Vertical profiles of particle-associated (>2.7 µm) and free-living (0.2-2.7 µm) functional gene concentrations of (a) RuBisCO-I large subunit (*cbbL*), representing eukaryote algae and/or cyanobacteria (b) RuBisCO-II small subunit (*cbbM*), representing purple sulfur bacteria and/or sulfur-oxidizing chemoautotrophic Gammaproteobacteria (c) ATP citrate lyase (*aclB*), representing green sulfur bacteria and/or sulfur-oxidizing chemoautotrophic Epsilonbacteraeota (c) methyl coenzyme reductase subunit A (*mcrA*), representing methanogens and/or anaerobic methane oxidizers collected in July 2017. Broken lines indicate oxycline boundaries. Error bars represent ± 1 S.D. of triplicate analyses.

## V-Discussion

To better understand how δ^13^C_OC_ of well-preserved Proterozoic sedimentary rock may reflect the past activities of autotrophic and methane-cycling populations, we quantified carbon fixation pathway gene copy numbers and measured δ^13^C_POC_ of size-fractionated particulate matter recovered from meromictic FGL. A size-fractionating approach was taken because biomass carbon associated with aggregation and particles is preferentially deposited in sediments through the biological pump and subsequently incorporated into sedimentary rock (Alldredge and Gotschalk, 1988, Shen et al., 2018). In FGL, we found that most of the primary production and biomass production occurs in the deeper, sulfidic photic zone and is primarily attributable to photoautotrophic anoxygenic purple sulfur bacteria in the larger size fraction. This particle-associated and/or aggregated biomass has a high probability of evading remineralization and being incorporated into the lakebed sediment for the following reasons. Anoxic conditions reduce the likelihood of ingestion by mesozooplankton. Additionally, as particle size and settling velocity increase, particle ingestion by mesozooplankton and protists becomes more problematic. Furthermore, high particle abundances in the light-scattering layer promote particle-particle collisions, enhance aggregation, and promote vertical transport (Burd and Jackson, 2009). Therefore, lakebed sediment may preferentially reflect input from purple sulfur bacteria layer.

Our qPCR (**Figure 7**), microscopy (**Error! Reference source not found.**), and field data (**Figure 1**, **Figure 2**) agree with our 2017 amplicon libraries, showing that the light-scattering layer consisted of three distinctive photoautotrophic populations that are physically partitioned by their light requirements and hydrogen sulfide tolerances (Cohen et al., 2023). Therefore, we expected shallower oxic and hypoxic photic zone δ^13^C_POC_ of both size fractions to reflect cyanobacterial biomass (high-O_2_ CBB), δ^13^C_POC_ of both size fractions in the deeper photic zone to reflect purple and green sulfur bacteria and thioautotrophic epsilonproteobacteria biomass (low-O_2_ CBB + rTCA), and the PA fraction δ^13^C_POC_ in the aphotic zone to reflect sinking purple sulfur bacteria. We did not expect methane cycling to strongly impact δ^13^C_POC_. Although maximum *mcrA* copy numbers in both size fractions occurred at the lower-oxycline boundary, the coincident highly depleted δ^13^C_POC_ in both size fractions cannot be attributed to abundant methanogen or methanotroph populations, because amplicons for only one methanogen, *Methanoregula*, were a minor contributor to 16S rRNA gene libraries (Cohen et al., 2023). Furthermore, no anaerobic methane-oxidizing taxa were recovered in these libraries and only modest aerobic methanotroph populations were restricted to oxic and hypoxic waters (Cohen et al., 2023).

The δ^13^C data suggest that all photoautotroph populations near the lower oxycline boundary assimilate isotopically-depleted bicarbonate entering the lake as groundwater. The admixture of groundwater with lake water results in δ^13^C_DIC_ values of -12.5‰ near the lower oxycline boundary (Havig et al., 2018, **Figure 6b**). Values reported in **Table 1** are collated from marine studies, and therefore assume an average δ^13^C_DIC_ of 1.7‰ in seawater (Cheng et al., 2019). Deriving approximate fractionation (ε) factors from **Table 1**, we estimate a δ^13^C_POC_ of ∼ -31.5 to -37.0‰ through low-O_2_ CBB by purple sulfur bacteria, δ^13^C_POC_ of ∼ -10.5 to -11.5‰ through rTCA by green sulfur bacteria, and δ^13^C_POC_ of ∼ -39.5 to -47.5‰ through high-O_2_ CBB by cyanobacteria, resulting in a three end-member average δ^13^C_POC_ near the lower oxycline of -27.2 to -32.0 ‰ assuming equal contribution from each, which is unlikely (**Figure 8**). Nevertheless, this range of approximate values is much heavier than our measured δ^13^C_POC_ -41.9‰, and -39.4 ‰ in PA and FL fractions at the lower oxycline boundary, where all three photoautotroph populations co-exist. The lower oxycline boundary is also where total particle concentrations (measured turbidity) peak, so that particle-particle collisions should also be highest and result in the most similarity between δ^13^C_POC_ values of the two size-fractions. As there are no aerobic methanotrophs at these depths, we suggest that the abundant sulfate-reducing bacteria, including the strongly particle-associated *Desulfocapsa* population at 20.5 m and the primarily free-living sulfate-reducing Deltaproteobacterial populations at 20.0 m (e.g., *Desulfatiglans, Syntrophus, Desulfobacca, Desulfovibrio*) (Cohen et al., 2023) may contribute a substantial amount of ^13^C-depleted biomass (**Figure 8**). The greatest abundance of *Desulfocapsa* 16S rRNA genes occurs at the same depth as the PA negative isotopic excursion, while the greatest abundance of the Deltaproteobacterial sulfate-reducers’ 16S rRNA genes occur at the same depth as the FL negative isotopic excursion, maximum measured bacterial heterotrophic production, and the first appearance of hydrogen sulfide (**Figure 1 Figure 4 Figure 6**). Bacterial heterotrophic production as measured in this study is often considered a proxy for protein remineralization, which may produce low molecular weight organic acids such as acetate, butyrate, and formate. The oxidation of these organic acids is paired with the reduction of sulfate to hydrogen sulfide by sulfate-reducers (Jørgensen et al., 2001). Therefore, oxidation of acetate by acetyl-CoA decarbonylase via the sulfate-reducing Woods Ljungdahl pathway by free-living sulfate-reducers is expected to be elevated at this depth.

**Figure 8:**
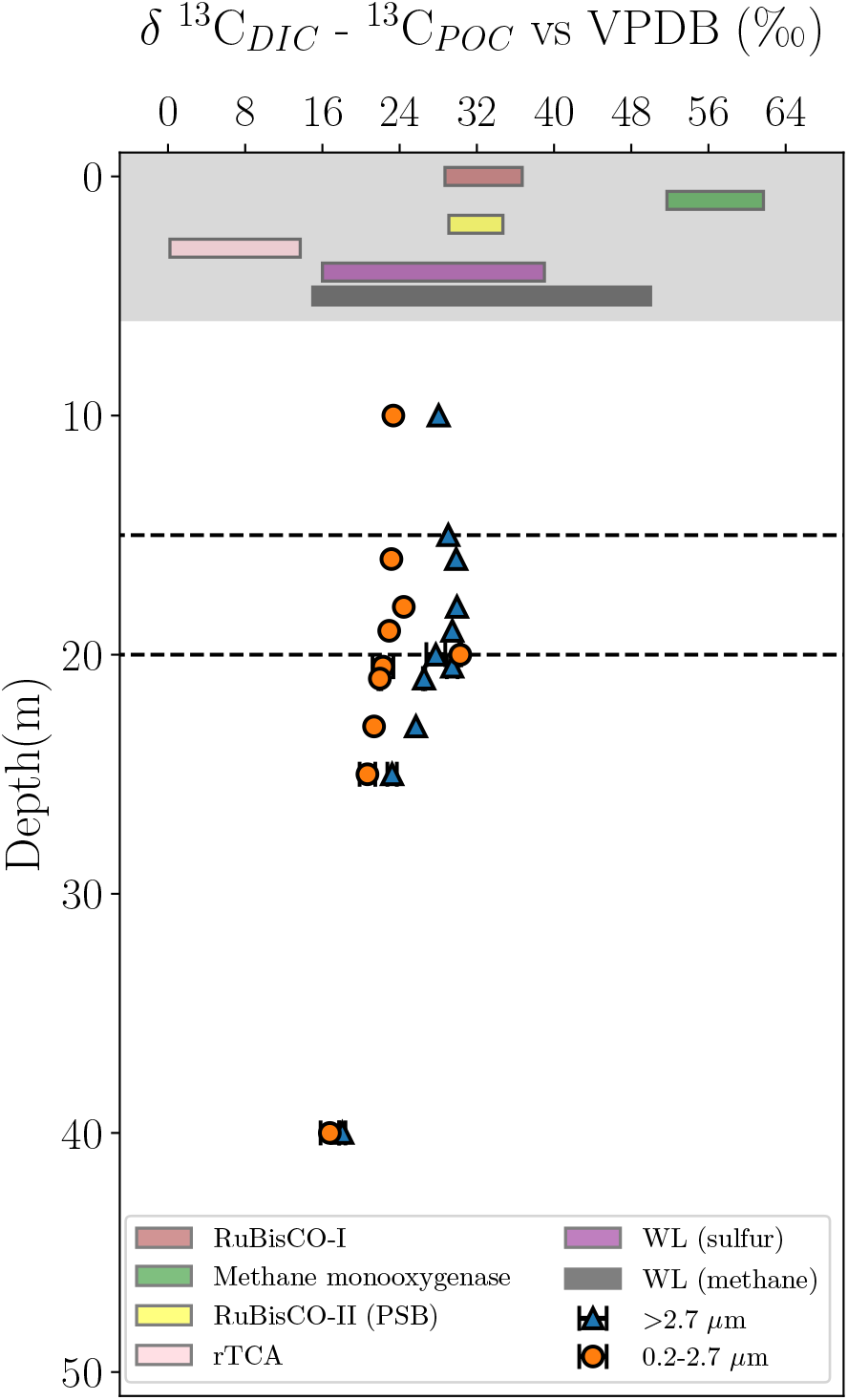
Depth profile of differences between Havig et al. (2018) dissolved inorganic carbon (DIC) and our July 2018 FL (0.2-2.7 µm and PA (>2.7 µm) size fraction particulate organic carbon (POC) stable isotopic compositions. These differences, approximate isotopic fractionation factors (ε), are compared to ranges of carbon fixation pathways’ ε. Pathway ranges are represented by colored bars in the shaded area and have no relationship to depth in this plot. Broken lines indicate oxycline boundaries. Error bars represent ± 1 S.D. of duplicate analyses. WL=Woods-Ljungdahl, PSB=Purple Sulfur Bacteria, rTCA=reverse citric acid cycle.

At 20.5 m, purple sulfur bacteria that likely contribute to the majority of light DIC assimilation (Culver and Brunskill, 1969) are the most abundant (Cohen et al., 2023) autotrophs. The δ^13^C_POC_ values of the PA fraction at this depth is therefore surprisingly depleted (−42.3‰), especially given that low-O_2_ CBB-indicating *cbbM* gene copy numbers (δ^13^C_cbbM_ values = -19 to -25 ‰, yielding ε of ý 20.7-26.7) were especially high in the PA fraction. The FL fraction δ^13^C_POC_ at this depth was far more ^13^C-enriched. This was expected because carbon assimilation via the rTCA pathway by green sulfur bacteria, which modestly contribute to the maximum light carbon assimilation rate (Culver and Brunskill, 1969) and chemoautotrophic epsilonproteobacteria, which are likely associated with the maximum dark carbon assimilation rate (Cohen et al., 2021), were dominant in this fraction. The most ^13^C-enriched δ^13^C_POC_ values observed in both size fractions of the turbidity layer (21 m) coincides with maxima in green sulfur bacteria and *aclB* gene abundances indicating that the rTCA fixation pathway dominated at that depth. Increasing δ^13^C_POC_ values with depth in the PA fraction and near-constant δ^13^C_POC_ values in the FL fraction in aphotic waters along with increases in PA *cbbM*, *aclB, and cbbL* copy numbers (*cbbM* ý *cbbL* > *aclB*) near the lakebed (40 m) support our hypothesis that a mixture of anoxygenic and oxygenic photoautotrophs contribute to the δ^13^C_POC_ values of lakebed organic matter, superficially resembling the δ^13^C_POC_ values of shallow-dwelling oxygenic algae (**Figure 8**). FGL lakebed sediments are classified as sapropel (1.8-2.4 weight % OC), with surface sediment δ^13^C_POC_ values being -32.6 ‰ (Havig et al., 2018). Given the δ^13^C_POC_ of our deepest (40 m) > 2.7 µm sample is - 37‰, and that the change in bulk δ^13^C_POC_ values over depth throughout the aphotic (below 23 m) monimolimnion appears to be linear (Fulton et al., 2018), we extrapolate to approximately the same value at the sediment-water interface. One possibility for this difference between sediment and water column δ^13^C values is further remineralization in the bottom 12 m of the water column and lakebed remineralization. Our hypothesis is supported by direct bright-field micrographs of 40 m samples containing mostly aggregates of purple sulfur bacteria with lesser amounts of cyanobacterial and green sulfur bacterial cells, and no eukaryote algae (**Figure S2b**).

## VI-Summary/Conclusion

To better understand how preserved ancient δ^13^C_OC_ values reflect past activities of autotrophs and methane-cyclers, we measured δ^13^C_POC_ values and carbon-fixation pathway marker genes of size-fractionated plankton corresponding to particle-associated/aggregated/symbiont and free-living microbes through all redox zones in the meromictic FGL. Size partitioning of carbon fixation genes and their vertical distributions reflected populations of photoautotrophs and methanogens. These include mostly free-living cyanobacteria in the shallow oxic and hypoxic photic zone (high oxygen-CBB, *cbbL* gene), mostly particle-associated purple sulfur bacteria (low-oxygen CBB, *cbbM* gene) and equally size-fractionated green sulfur bacteria (reverse citric acid cycle, *aclB* gene) in the euxinic photic zone, and equally size-fractionated methanogens (Wood-Ljungdahl pathway, *mcrA* gene) at the lower oxycline boundary. The δ^13^C values of lakebed sediments taken at face value might be interpreted as being derived from shallow eukaryote algae or cyanobacteria. However, our results show that δ^13^C values of particles arriving at the lakebed reflect a mixture of populations exported from the particle-rich (turbid) hypoxic and euxinic deep photic zone that assimilate a groundwater-derived, isotopically depleted inorganic carbon source or pair sulfate reduction with volatile fatty acid oxidation. This suggests that organic-rich Proterozoic sediments deposited in waters with a sulfidic photic zone could reflect a mixture of carbon fixation pathways that superficially resembles the δ^13^C_POC_ of eukaryote algae in the oxygenated photic zone.

**Data availability:** Data is available as downloadable supplementary tables.

## Supporting information

qPCR

biological_metadata

chemical_metadata

isotope_data

## References

1. Alldredge, A.L. and Gotschalk, C. 1988. In situ settling behavior of marine snow 1. Limnology and Oceanography, 33(3): 339–351. doi: 10.4319/lo.1988.33.3.0339

2. Anbar, A.D. and Knoll, A.H. 2002. Proterozoic ocean chemistry and evolution: a bioinorganic bridge? Science. 297(5584): 1137–1142. doi: 10.1126/science.1069651

3. Berg, I.A. 2011. Ecological aspects of the distribution of different autotrophic CO2 fixation pathways. Applied and environmental microbiology. 77(6):1925–1936. doi: 10.1128/AEM.02473-10

4. Blotta, I., Prestinaci, F., Mirante, S. and Cantafora, A. 2005. Quantitative assay of total dsDNA with PicoGreen reagent and real-time fluorescent detection. Annali-Istituto Superiore DiSanita, 41(1): 119.

5. Brunskill, G.J. and Ludlam, S.D. 1969. Fayetteville Green Lake, New York. I. Physical and chemical limnology 1. Limnology and Oceanography. 14(6): 817–829. doi: 10.4319/lo.1969.14.6.0817

6. Burd, A.B. and Jackson, G.A. 2009. Particle aggregation. Annual review of marine science. 1: 65–90. doi: 10.1146/annurev.marine.010908.163904

7. Campbell, B.J., Stein, J.L. and Cary, S.C. 2003. Evidence of chemolithoautotrophy in the bacterial community associated with Alvinella pompejana, a hydrothermal vent polychaete. Applied and environmental microbiology. 69(9): 5070–5078. doi: 10.1128/AEM.69.9.5070-5078.2003

8. Campbell, B.J. and Cary, S.C. 2004. Abundance of reverse tricarboxylic acid cycle genes in free-living microorganisms at deep-sea hydrothermal vents. Applied and environmental microbiology. 70(10): 6282–6289. doi: 10.1128/AEM.70.10.6282-6289.2004

9. Canfield, D.E., Stewart, F.J., Thamdrup, B., De Brabandere, L., Dalsgaard, T., Delong, E.F., Revsbech, N.P. and Ulloa, O. 2010. A cryptic sulfur cycle in oxygen-minimum–zone waters off the Chilean coast. Science. 330(6009): 1375–1378. doi:10.1126/science.1196889

10. Cheng, L., Normandeau, C., Bowden, R., Doucett, R., Gallagher, B., Gillikin, D.P., Kumamoto, Y., McKay, J.L., Middlestead, P., Ninnemann, U. and Nothaft, D. 2019. An international intercomparison of stable carbon isotope composition measurements of dissolved inorganic carbon in seawater. Limnol. Oceanog. Methods: 17(3): 200–209. doi: 10.1002/lom3.10300

11. Cohen, A. B., Novkov-Bloom, A., Wesselborg, C., Yagudaeva, M., Aranguiz, E., and Taylor, G. T. 2021. Applying fluorescence in situ hybridization to aquatic systems with cyanobacteria blooms: Autofluorescence suppression and high-throughput image analysis. Limnology and Oceanography Methods. 19: 457–475. doi: 10.1002/lom3.10437

12. Cohen, A.B., Klepac-Ceraj, V., Bidas, K., Weber, F., Garber, A.I., Christensen, L.N., Cram, J.A., McCormick, M.L. and Taylor, G.T. 2023. Deep photoautotrophic prokaryotes contribute substantially to carbon dynamics in oxygen-deficient waters in a permanently redox-stratified freshwater lake. Limnol. Oceanog. 68(1): 232–247. doi: 10.1002/lno.12262

13. Culver, D.A. and Brunskill, G.J. 1969. Fayetteville Green Lake, New York. V. Studies of primary production and zooplankton in a meromictic marl lake 1. Limnol. Oceanog. 14(6): 862–873. doi: 10.4319/lo.1969.14.6.0862

14. Des Marais, D.J. 1997. Isotopic evolution of the biogeochemical carbon cycle during the Proterozoic Eon. Organic Geochemistry. 27(5-6): 185–193. doi: 10.1016/S0146-6380(97)00061-2

15. Denman, S.E., Tomkins, N.W. and McSweeney, C.S. 2007. Quantitation and diversity analysis of ruminal methanogenic populations in response to the antimethanogenic compound bromochloromethane. FEMS microbiology ecology. 62(3):313–322. doi: 10.1111/j.1574-6941.2007.00394.xx

16. Dhillon, A., Lever, M., Lloyd, K.G., Albert, D.B., Sogin, M.L. and Teske, A. 2005. Methanogen diversity evidenced by molecular characterization of methyl coenzyme M reductase A (mcrA) genes in hydrothermal sediments of the Guaymas Basin. Applied and Environmental Microbiology. 71(8): 4592–4601. doi: 10.1128/AEM.71.8.4592-4601.2005

17. Eigenbrode, J.L. and Freeman, K.H. 2006. Late Archean rise of aerobic microbial ecosystems. Proceedings of the National Academy of Sciences. 103(43): 15759–15764. doi: 10.1073/pnas.060754010

18. Emerson, S. and Hedges, J. 2008. Chemical oceanography and the marine carbon cycle. Cambridge University Press.

19. Fulton, J.M., Arthur, M.A., Thomas, B. and Freeman, K.H. 2018. Pigment carbon and nitrogen isotopic signatures in euxinic basins. Geobiology. 16(4): 429–445. doi:10.1111/gbi.12285

20. Hallam, S.J., Putnam, N., Preston, C.M., Detter, J.C., Rokhsar, D., Richardson, P.M. and DeLong, E.F. 2004. Reverse methanogenesis: testing the hypothesis with environmental genomics. Science. 305(5689): 1457-1462. doi: 10.1126/science.1100025

21. Hanson, R.S. and Hanson, T.E. 1996. Methanotrophic bacteria. Microbiological reviews. 60(2): 439–471.

22. Havig, J.R., Hamilton, T.L., McCormick, M., McClure, B., Sowers, T., Wegter, B. and Kump, L.R. 2018. Water column and sediment stable carbon isotope biogeochemistry of permanently redox-stratified Fayetteville Green Lake, New York, USA. Limnology and Oceanography.63(2): 570–587. doi: 10.1002/lno.10649

23. Hayes, J.M. 1993. Factors controlling 13C contents of sedimentary organic compounds: principles and evidence. Marine Geology. 113(1-2): 111–125. doi: 10.1016/0025-3227(93)90153-M

24. Hayes, J.M., Strauss, H. and Kaufman, A.J. 1999. The abundance of 13C in marine organic matter and isotopic fractionation in the global biogeochemical cycle of carbon during the past 800 Ma. Chemical Geology. 161(1-3): 103–125. doi: 10.1016/S0009-2541(99)00083-2.

25. Hayes, J.M. 2001. Fractionation of carbon and hydrogen isotopes in biosynthetic processes. Reviews in mineralogy and geochemistry. 43(1). 225–277. doi: 10.2138/gsrmg.43.1.225

26. Hügler, M. and Sievert, S.M. 2011. Beyond the Calvin cycle: autotrophic carbon fixation in the ocean. Annual review of marine science. 3: 261–289. doi: 10.1146/annurev-marine-120709-142712

27. Imhoff, J.F., 1995. Taxonomy and physiology of phototrophic purple bacteria and green sulfur bacteria. Anoxygenic photosynthetic bacteria: 1–15.

28. Grote, J., Jost, G., Labrenz, M., Herndl, G.J. and Jürgens, K. 2008. Epsilonproteobacteria represent the major portion of chemoautotrophic bacteria in sulfidic waters of pelagic redoxclines of the Baltic and Black Seas. Applied and environmental microbiology 74(24): 7546–7551. doi: 10.1128/AEM.01186-08

29. Johnston, D.T., Wolfe-Simon, F., Pearson, A. and Knoll, A.H. 2009. Anoxygenic photosynthesis modulated Proterozoic oxygen and sustained Earth’s middle age. Proceedings of the National Academy of Sciences. 106(40): 16925–16929. doi: 10.1073/pnas.0909248106

30. Jørgensen, B.B., Weber, A. and Zopfi, J. 2001. Sulfate reduction and anaerobic methane oxidation in Black Sea sediments. Deep Sea Research Part I: Oceanographic Research Papers. 48(9): 2097–2120. doi: 10.1016/S0967-0637(01)00007-3

31. Kump, L.R. and Arthur, M.A. 1999. Interpreting carbon-isotope excursions: carbonates and organic matter. Chemical Geology. 161(1-3): 181–198. doi: 10.1016/S0009-2541(99)00086-8

32. Lin, X., Wakeham, S.G., Putnam, I.F., Astor, Y.M., Scranton, M.I., Chistoserdov, A.Y. and Taylor, G.T. 2006. Comparison of vertical distributions of prokaryotic assemblages in the anoxic Cariaco Basin and Black Sea by use of fluorescence in situ hybridization. Applied and Environmental Microbiology. 72(4): 2679–2690. doi: 10.1128/AEM.72.4.2679-2690.2006

33. Lücker, S., Nowka, B., Rattei, T., Spieck, E., and Daims, H. 2013. The genome of Nitrospina gracilis illuminates the metabolism and evolution of the major marine nitrite oxidizer. Front Microbiol 4: 27. doi: 10.3389/fmicb.2013.00027

34. Luton, P.E., Wayne, J.M., Sharp, R.J. and Riley, P.W., 2002. The mcrA gene as an alternative to 16S rRNA in the phylogenetic analysis of methanogen populations in landfill. Microbiology. 148(11):.3521–3530. doi: 10.1099/00221287-148-11-3521

35. McDonald, I.R., Kenna, E.M. and Murrell, J.C. 1995. Detection of methanotrophic bacteria in environmental samples with the PCR. Applied and Environmental Microbiology.61(1):116–121.

36. McNevin, D.B., Badger, M.R., Whitney, S.M., Von Caemmerer, S., Tcherkez, G.G. and Farquhar, G.D. 2007. Differences in carbon isotope discrimination of three variants of D-ribulose-1, 5-bisphosphate carboxylase/oxygenase reflect differences in their catalytic mechanisms. Journal of Biological Chemistry. 282(49): 36068–36076.

37. Minamoto, T., Naka, T., Moji, K. and Maruyama, A. 2016. Techniques for the practical collection of environmental DNA: filter selection, preservation, and extraction. Limnology. 17: 23–32. doi: 10.1007/s10201-015-0457-4

38. Momper, L., Jungbluth, S.P., Lee, M.D. and Amend, J.P. 2017. Energy and carbon metabolisms in a deep terrestrial subsurface fluid microbial community. The ISME journal. 11(10): 2319–2333. doi: 10.1038/ismej.2017.94

39. Nunoura, T., Chikaraishi, Y., Izaki, R., Suwa, T., Sato, T., Harada, T., Mori, K., Kato, Y., Miyazaki, M., Shimamura, S. and Yanagawa, K. 2018. A primordial and reversible TCA cycle in a facultatively chemolithoautotrophic thermophile. Science. 359(6375): 559–563. doi: 10.1126/science.aao3407

40. Orphan, V.J., House, C.H., Hinrichs, K.U., McKeegan, K.D. and DeLong, E.F. 2001. Methane-consuming archaea revealed by directly coupled isotopic and phylogenetic analysis. Science. 293(5529):484–487. doi: 10.1126/science.1061338

41. Posth, N.R., Bristow, L.A., Cox, R.P., Habicht, K.S., Danza, F., Tonolla, M., Frigaard, N.U. and Canfield, D.E. 2017. Carbon isotope fractionation by anoxygenic phototrophic bacteria in euxinic Lake Cadagno. Geobiology. 15(6): 798–816. doi: 10.1111/gbi.12254

42. Probst, A.J., Castelle, C.J., Singh, A., Brown, C.T., Anantharaman, K., Sharon, I., Hug, L.A., Burstein, D., Emerson, J.B., Thomas, B.C. and Banfield, J.F. 2017. Genomic resolution of a cold subsurface aquifer community provides metabolic insights for novel microbes adapted to high CO2 concentrations. Environmental microbiology. 19(2): 459–474. doi: 10.1111/1462-2920.13362

43. Robinson, J.J. and Cavanaugh, C.M., 1995. Expression of form I and form II RuBisCO in chemoautotrophic symbioses: implications for the interpretation of stable carbon isotope values. Limnology and Oceanography. 40(8):1496–1502. doi: 10.4319/lo.1995.40.8.1496

44. Ruiz-Fernández, P., Ramírez-Flandes, S., Rodríguez-León, E. and Ulloa, O. 2020. Autotrophic carbon fixation pathways along the redox gradient in oxygen-depleted oceanic waters. Environmental microbiology reports. 12(3): 334–341. doi: 10.1111/1758-2229.12837

45. Shen, J., Pearson, A., Henkes, G.A., Zhang, Y.G., Chen, K., Li, D., Wankel, S.D., Finney, S.C. and Shen, Y. 2018. Improved efficiency of the biological pump as a trigger for the Late Ordovician glaciation. Nature Geoscience.11(7): 510–514. doi: 10.1038/s41561-018-0141-5

46. Tabita, F.R., Satagopan, S., Hanson, T.E., Kreel, N.E. and Scott, S.S. 2008. Distinct form I, II, III, and IV RuBisCO proteins from the three kingdoms of life provide clues about RuBisCO evolution and structure/function relationships. Journal of experimental botany. 59(7): 1515–1524. doi: 10.1098/rstb.2008.0023

47. Templeton, A.S., Chu, K.H., Alvarez-Cohen, L. and Conrad, M.E. 2006. Variable carbon isotope fractionation expressed by aerobic CH4-oxidizing bacteria. Geochimica et Cosmochimica Acta. 70(7): 1739–1752. doi: 10.1016/j.gca.2005.12.002

48. Thomazo, C., Pinti, D.L., Busigny, V., Ader, M., Hashizume, K. and Philippot, P. 2009. Biological activity and the Earth’s surface evolution: insights from carbon, sulfur, nitrogen and iron stable isotopes in the rock record. Comptes Rendus Palevol. 8(7): 665–678. doi: 10.1016/j.crpv.2009.02.003

49. Thompson, J.B., Ferris, F.G. and Smith, D.A. 1990. Geomicrobiology and sedimentology of the mixolimnion and chemocline in Fayetteville Green Lake, New York. Palaios. 5(1): 52–75.

50. Tolli, J. and King, G.M. 2005. Diversity and structure of bacterial Chemolithotrophic communities in pine forest and agroecosystem soils. Applied and Environmental Microbiology. 71(12): 8411–8418. doi: 10.1128/AEM.71.12.8411-8418.2005

51. Torgersen, T., Hammond, D.E., Clarke, W.B. and Peng, T.H. 1981. Fayetteville, Green Lake, New York: 3H–3He water mass ages and secondary chemical structure 1, 2. Limnology and Oceanography. 26(1): 110-122. doi: 10.4319/lo.1981.26.1.0110

52. Wahlund, T.M. and Tabita, F.R. 1997. The reductive tricarboxylic acid cycle of carbon dioxide assimilation: initial studies and purification of ATP-citrate lyase from the green sulfur bacterium Chlorobium tepidum. Journal of bacteriology. 179(15): 4859–4867. doi: 10.1128/jb.179.15.4859-4867.1997

53. Werne, J.P. and Hollander, D.J. 2004. Balancing supply and demand: controls on carbon isotope fractionation in the Cariaco Basin (Venezuela) Younger Dryas to present. Marine Chemistry. 92(1-4): 275–293. doi: 10.1016/j.marchem.2004.06.031

54. Zerkle, A.L., Kamyshny Jr, A., Kump, L.R., Farquhar, J., Oduro, H. and Arthur, M.A. 2010. Sulfur cycling in a stratified euxinic lake with moderately high sulfate: constraints from quadruple S isotopes. Geochimica et Cosmochimica Acta. 74(17): 4953–4970. doi: 10.1016/j.gca.2010.06.015

