## supplemental_text for "Preserved particulate organic carbon is likely derived from the subsurface sulfidic photic zone of the Proterozoic Ocean: evidence from a modern, oxygen-deficient lake"

Supplementary Materials

**TABLE S1**: Polymerase chain reaction tests for the presence or absence of the *mcrA* gene using Lucigen FailSafe™ reagents and primers in Luton et al. (2002). Dilution indicates fold-dilution with nuclease-free water. "+" indicates presence and "-" indicates absence. Reaction chemistry and thermal program are presented in **Table 2**.

| **Free-living** | *Sample ID* | *Depth (m)* | *dilution* | *mcrA* |
|---|---|---|---|---|
| | 102 | 10 | 1 | - |
| | | | 10 | - |
| | | | 100 | - |
| | 71 | 15 | 1 | - |
| | | | 10 | - |
| | | | 100 | - |
| | 73 | 16 | 1 | + |
| | | | 10 | - |
| | | | 100 | - |
| | 69 | 19 | 1 | - |
| | | | 10 | - |
| | | | 100 | - |
| | 55 | 19.75 | 1 | - |
| | | | 10 | - |
| | | | 100 | - |
| | 59 | 20 | 1 | - |
| | | | 10 | - |
| | | | 100 | + |
| | 54 | 20.5 | 1 | - |
| | | | 10 | - |
| | | | 100 | + |
| | 60 | 21 | 1 | - |
| | | | 10 | - |
| | | | 100 | + |
| | 50 | 23 | 1 | - |
| | | | 10 | + |
| | | | 100 | + |
| | 61 | 25 | 1 | - |
| | | | 10 | + |
| | | | 100 | + |
| | 70 | 40 | 1 | - |

| | | | | |
|---|---|---|---|---|
| | | | 10 | - |
| | | | 100 | - |
| **Particle-associated** | | | | |
| | *Sample ID* | *Depth (m)* | *dilution* | *mcrA* |
| | 102C | 10 | 1 | - |
| | | | 10 | - |
| | | | 100 | - |
| | 71C | 15 | 1 | - |
| | | | 10 | - |
| | | | 100 | - |
| | 73C | 16 | 1 | + |
| | | | 10 | - |
| | | | 100 | - |
| | 69C | 19 | 1 | |
| | | | 10 | - |
| | | | 100 | |
| | 55C | 19.75 | 1 | |
| | | | 10 | |
| | | | 100 | |
| | 59C | 20 | 1 | |
| | | | 10 | |
| | | | 100 | + |
| | 54C | 20.5 | 1 | |
| | | | 10 | |
| | | | 100 | + |
| | 60C | 21 | 1 | |
| | | | 10 | |
| | | | 100 | + |
| | 50C | 23 | 1 | - |
| | | | 10 | - |
| | | | 100 | + |
| | 61C | 25 | 1 | - |
| | | | 10 | + |
| | | | 100 | + |
| | 70C | 40 | 1 | - |
| | | | 10 | - |
| | | | 100 | - |

(a) 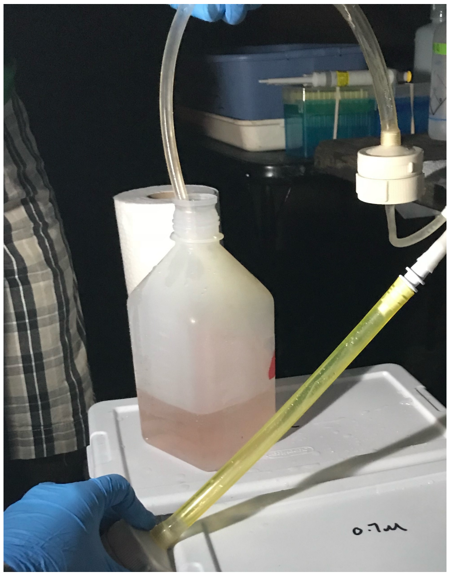 (b) 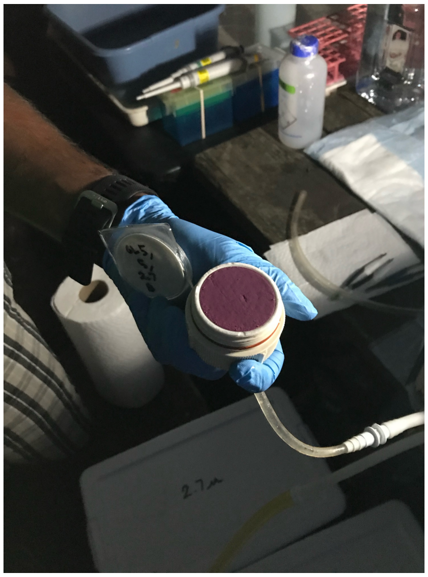

**Figure S1:** Water recovered from 20.5m depth in Fayetteville Green Lake during July 2018 was visibly purple (a), and coloration was from high concentrations of > 2.7 µm purple sulfur bacteria aggregates (b).

(a) 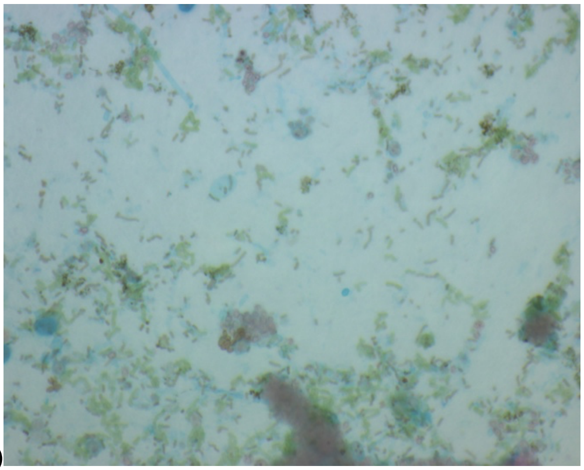 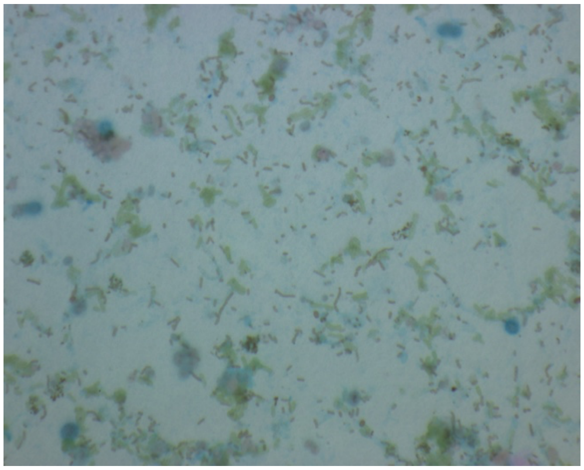

(b) 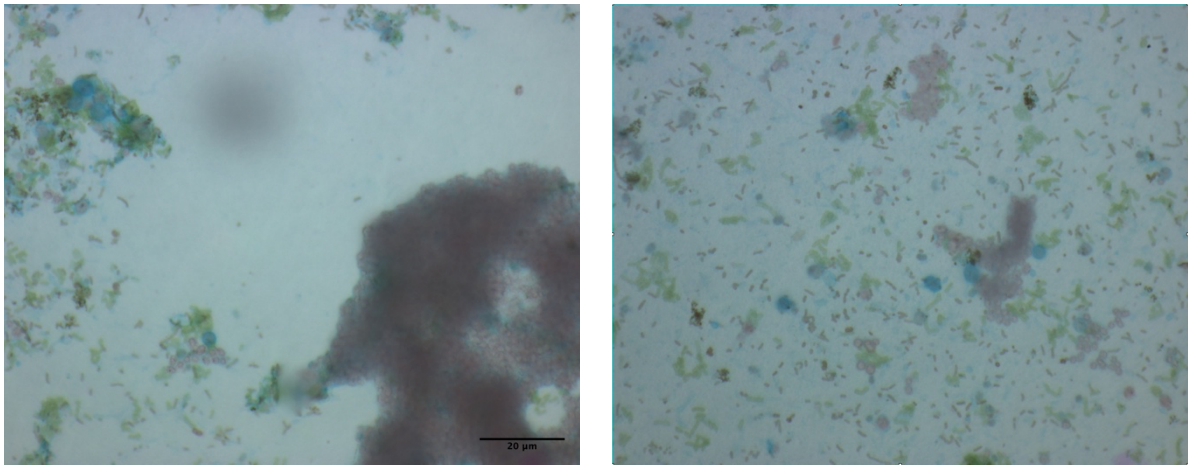


**Figure S2:** Light micrographs of Fayetteville Green Lake water recovered from (a) 21 m and (b) 40 m in July 2017 and stained with transparent exopolymeric substance-indicating Alcian Blue dye as in Engel, 2009. Micrographs in (a) show that green sulfur bacteria are the dominant photoautotroph at 21 m. Micrographs in (b) illustrate the presence of large purple sulfur bacteria aggregates and smaller green sulfur bacteria and cyanobacteria interspersed with the transparent exopolymeric substances derived from the light scattering layer (19-23) in the deep lake. Micrographs were taken at 630x magnification using a Zeiss Axioscope microscope coupled to an Optronics MagnaFire™ CCD camera.

**References**


Engel, A., 2009. Determination of marine gel particles. In Practical guidelines for the analysis of seawater: 125-142.

Luton, P.E., Wayne, J.M., Sharp, R.J. and Riley, P.W., 2002. The mcrA gene as an alternative to 16S rRNA in the phylogenetic analysis of methanogen populations in landfill. Microbiology. 148(11):.3521-3530. doi: 10.1099/00221287-148-11-3521
